# Systematic assessment of machine learning-based variant annotation methods for rare variant association testing

**DOI:** 10.64898/2026.03.18.712715

**Authors:** Matthew Aguirre, Flaviyan Jerome Irudayanathan, Maggie Crow, Hussein A. Hejase, Vipin K. Menon, Rion K. Pendergrass, Mark I. McCarthy, Kipper Fletez-Brant

## Abstract

Machine learning-based annotation methods are increasingly used to assess the pathogenicity of genetic variants, but their performance at prioritizing variants for gene-level association testing remains poorly characterized. Here, we systematically benchmark five annotation methods — CADD v1.6, CADD v1.7, AlphaMissense, ESM-1b, and GPN-MSA — using four primary gene-based tests and six annotation-level aggregation tests across 14 quantitative traits measured in up to 350,377 UK Biobank participants. Using a novel framework based on Wasserstein distances, we quantify how annotation choice affects test calibration and power. Tests using CADD annotations achieve the highest signal separation, while tests using AlphaMissense annotations exhibit systematically lower calibration. All combinations of methods produced significant results that were enriched (1.8–5.8-fold) for loss-of-function intolerant genes, though tests using GPN-MSA annotations displayed the highest such enrichment. Replication across symmetric phenotypes and loss-of-function burden tests was generally similar across methods. Our analysis provides practical guidance for annotation method selection in rare variant studies and establishes a distributional framework for calibration assessment.

## Introduction

As cohorts with linked genetic and clinical data have reached biobank scale, rare variant association tests (RVATs) are emerging as a powerful approach to estimate the effects of genes on complex traits [1, 2, 3]. Discoveries made using these tests are often biologically interpretable and complementary to signal from genome-wide association studies (GWAS) of common variants [4, 5]. However, the success of RVATs depend on the inclusion criteria used to define variant sets for testing. These criteria are often simple filters of variants according to their allele frequency and predicted functional consequences — most commonly, gene loss of function [1, 2, 3].

Meanwhile, machine learning-based variant scoring methods have gained popularity for clinical variant prioritization and also have potential for use in defining variant masks for gene aggregation tests [6, 7]. These scoring methods range from traditional ensemble models like CADD [8, 9], which combine multiple genomic annotations, to sequence-based deep learning models like AlphaMissense, ESM-1b, and GPN-MSA [10, 11, 12]. While these methods show promise for pathogenicity prediction, performing well on benchmarks using reference datasets like ClinVar [10, 12, 13, 14], their performance at nominating variants for inclusion in association tests remains underexplored. At the same time, the power of aggregation tests for missense and loss-of-function variants, including tests using deleteriousness labels from AlphaMissense, has been shown to vary [7, 15, 16].

Here, we present a systematic benchmark of five annotation methods — CADD v1.6 [8], CADD v1.7 [9], AlphaMissense [10], ESM-1b [11], and GPN-MSA [12] — as applied to 9,335,541 coding variants for use in ten gene-level rare variant association tests [17, 18, 19, 20, 21, 22, 23]. We analyze data on 14 quantitative traits in up to 350,377 UK Biobank participants [24], assessing performance using measures of genomic inflation [25] and a novel distributional framework based on Wasserstein distances. We use this approach to characterize trade-offs between test calibration and discovery power, and further describe how the genes prioritized using these methods capture signal across the functionally constrained genome. Our results offer guidance for annotation method selection in rare variant association studies and establish a robust framework for evaluating the calibration properties of association tests.

## Results

### Annotation methods differ in variant classification

We applied five commonly used variant annotation methods — CADD v1.6 [8], CADD v1.7 [9], AlphaMissense [10], ESM-1b [11], and GPN-MSA [12] — to all 6,218,579 missense and 3,116,962 synonymous coding variants called in gnomAD v4.1 [15] (9,335,541 protein coding variants in total, separately counting each alternate for multi-allelic variants; **Methods**). We then labeled each variant as benign, moderate, or deleterious using method-specific thresholds previously established in the literature (**Methods**).

Annotation methods showed pronounced differences in the proportion of variants classified as benign, moderate, or deleterious, and in the overall distribution of scores across variant types (**Fig. 1a**, **Table S1**, **Fig. S1**; **Methods**). Both CADD versions had permissive deleteriousness filters while ESM-1b and AlphaMissense applied more stringent criteria, and only a small fraction of missense variants (8.9%, 555,001 in total) were labeled as deleterious by all five methods (**Fig. 1a,b**). ESM-1b and GPN assigned the most variants to the moderate category (**Fig. 1a**), with broad agreement in the set of missense variants labeled as such (**Fig. S2**). Despite this overall divergence in labeling schemes, the raw pathogenicity scores for each variant were highly correlated by rank across methods (missense variants in **Fig. 1c**; results for all variants shown in **Fig. S4**). Relatedly, over 50% of benign-classified variants were labeled as such by all five methods, suggesting a consensus “null” set consisting mainly of synonymous variants (1,562,505 synonymous and 167,453 missense variants are in this consensus set of 1,729,958 in total; **Fig. 1d**; **Methods**). However, benign labels were generally discordant for missense variants, suggesting most overlap in this label set is due to agreement on synonymous variants (**Fig. S3**).

**Fig. 1:**
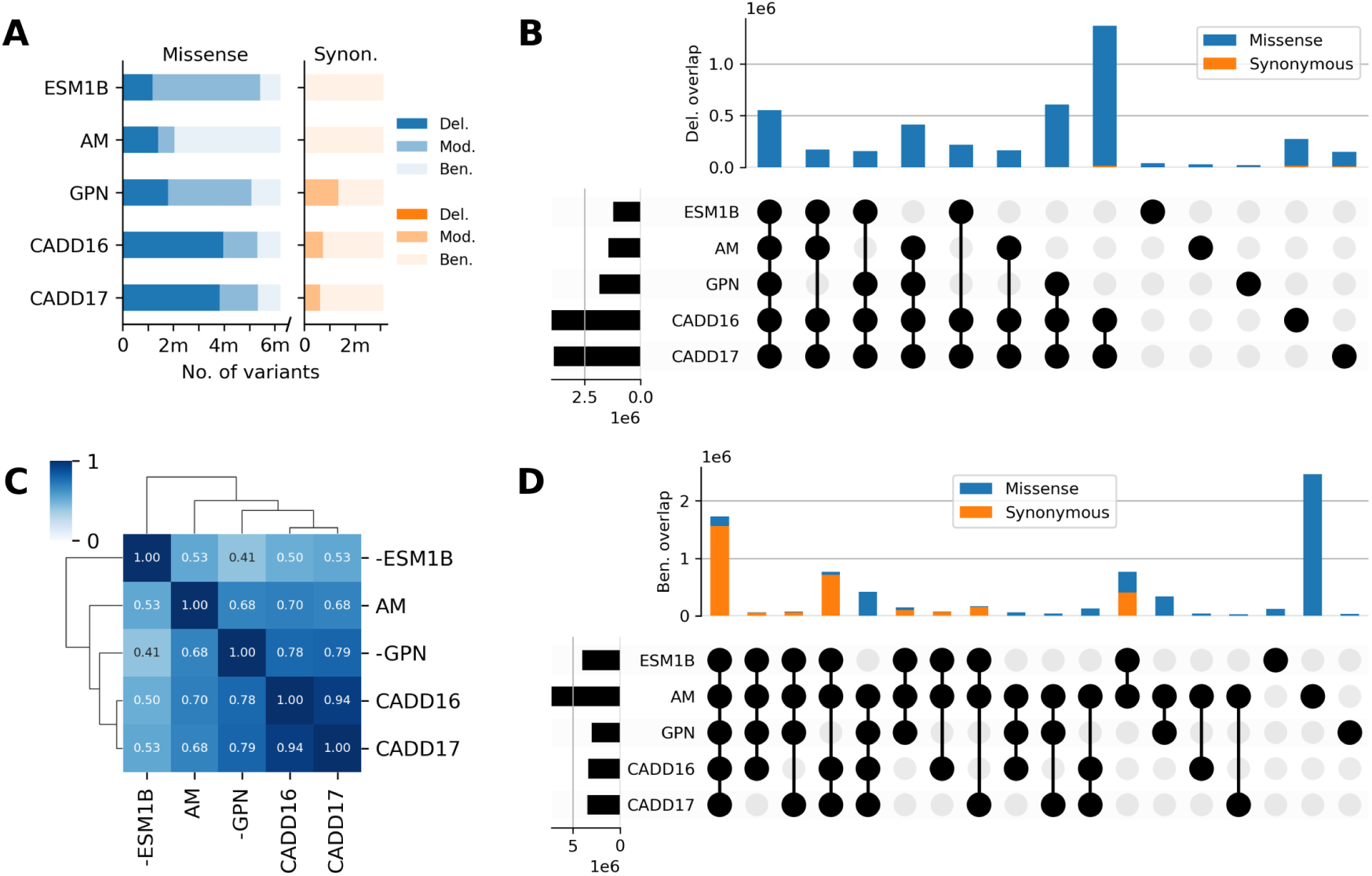
Variant annotation overlap. **(A)** Proportion of variants labeled by each annotation method as deleterious (dark red), moderate (medium red), and benign (light red), stratified by the variant’s most severe consequence on a gene (missense, on the left, or synonymous, on the right). **(B)** UpSet plot showing overlap in variants labeled as deleterious by each of the methods. The distribution of 43,894 synonymous variants is overwhelmed by that of 4,240,678 missense variants in this panel. Bars are not shown if they account for less than 0.2% of the total variant set. **(C)** Clustered heatmap of raw scores from the methods for missense variants. Numbers in each box indicate the rank correlation (Spearman *ρ*) between two methods. Scores from ESM and GPN are negated to polarize scores from all methods to the same direction (larger is more deleterious). **(D)** UpSet plot showing overlap in variants labeled as benign by each of the methods. There are 3,116,962 synonymous and 4,367,688 missense variants in total in this panel. Bars are not shown if they account for less than 0.2% of the total variant set.

### Genomic inflation varies by annotation method and statistical test

As an initial assessment of the calibration of tests using these labeled variant sets, we computed genomic inflation factors (*λ*_GC_) on gene *p*-values from non-common variants (MAF *<* 1%; see **Methods**) that were labeled as benign by each annotation method, under the assumption that these variants should tend to have minimal trait effects and therefore follow a distribution close to the null. However, we observed systematic differences in genomic inflation across the four primary rare variant aggregation tests (BURDEN, ACAT-V, SKAT, and SKAT-O [17, 18, 19]; **Fig. 2**) and six secondary tests that further combine information across annotation labels (BURDEN-ACAT, ACAT-V-ACAT, SKAT-O-ACAT, SBAT, GENE_P, and COAST-O [23, 21, 22]; **Fig. S5**), in cohorts of related and unrelated individuals (**Methods**; results including both related and unrelated individuals in **Fig. S7**).

**Fig. 2:**
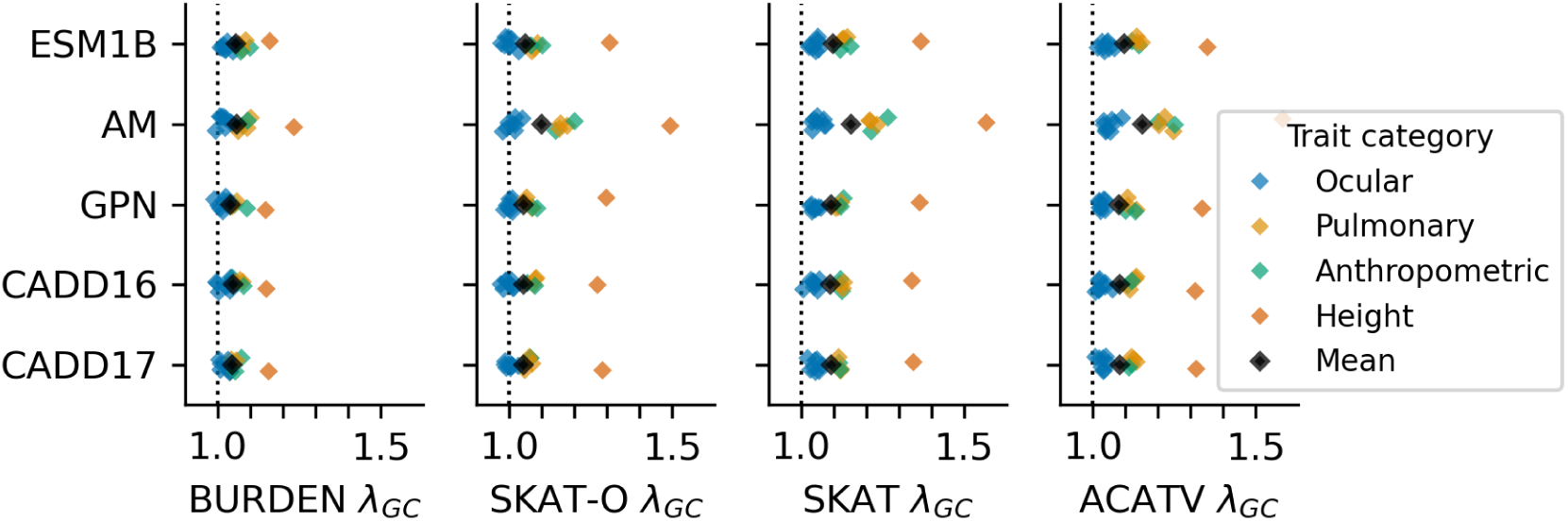
Genomic inflation by primary test and annotation method. Genomic inflation (*λ*_GC_) of primary test methods (subpanels) using variants classified as benign by each of the anno-tation methods (*y*-axis bins). Each point corresponds to one of 14 traits × 5 annotations × 4 tests = 280 RVATs conducted in a cohort of unrelated individuals, and is colored by trait category. Black points are the mean trait *λ*_GC_ for each combination of association test and annotation method.

Among primary test methods, burden and hybrid (SKAT-O) tests showed the best calibration (mean *λ*_GC_ = 1.04–1.12 across the five annotation methods; black points in **Fig. 2**), while the other variance component tests (SKAT and ACAT-V) exhibited slightly higher inflation (mean *λ*_GC_ = 1.05–1.17). Tests using AlphaMissense consistently produced the highest inflation across traits and test cohorts (*λ*_GC_ up to 1.8 for ACAT-V on height in the “all” cohort; **Fig. S7**), while on balance both CADD versions and GPN-MSA maintained the lowest average inflation (*λ*_GC_ ≈ 1.07; average across panels in **Fig. 2**). Three-way ANOVA confirmed that annotation method (*F* = 5.6*, p* = 2.2 × 10^-4^) and statistical test (*F* = 11.8*, p* = 1.8 × 10^-7^) significantly influenced genomic inflation, while test cohort did not have a significant effect (**Table S2**). Pairwise comparisons using Tukey’s HSD test showed that these differences were due to superior inflation control by SKAT-O and the burden test, and higher inflation in tests using AlphaMissense variant masks (**Tables S3, S4**).

Among secondary tests, which aggregate signal across the variant sets produced by each annotation method, we computed genomic inflation factors for the final set of test statistics for each gene (**Fig. S5**). Unlike for primary tests, we find no effect of annotation method on test statistic inflation (*p* = 0.91): differences are entirely driven by statistical test (*p* = 1.6 × 10*^−^*^69^) and cohort composition (*p* = 1.4 × 10*^−^*^3^; all results from three-way ANOVA, **Table S5**), reflecting that these tests efficiently capture signal from all scored variants when aggregating over annotation label sets. On balance, the allelic series and aggregated burden tests had lower inflation than the aggregated variance component tests, with GENE_P instead exhibiting systematic deflation (**Table S6**) and results overall mirroring trends in primary test inflation.

### Differential power and calibration across annotation method and statistical tests

To move beyond the point estimates of calibration provided by genomic inflation, we developed a distributional framework based on optimal transport to assess null calibration and power using the 1-Wasserstein (*W*_1_) distance. Rather than looking at a single quantile of association test results (e.g., using the median *χ*^2^ statistic to define *λ*_GC_), we measured the differences in distribution between tests that use benign and deleterious variant masks (**Fig. 3**). For each combination of annotation method and primary association test, we quantified calibration error as *W*_1_ between empirical benign-mask *χ*^2^ statistics and the theoretical 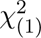 distribution, and signal separation as *W*_1_ between benign and deleterious mask *χ*^2^ statistics (**Methods**; see F**ig. S6** for a further explanatory diagram).

**Fig. 3:**
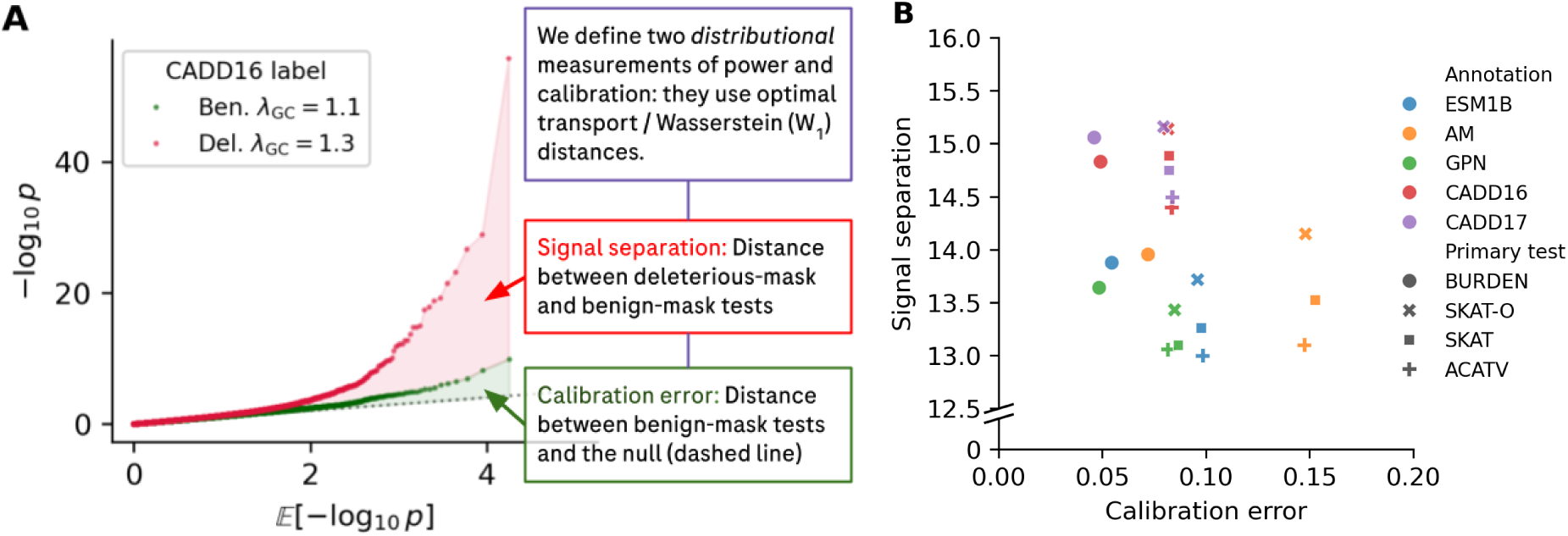
Distributional measures of power and calibration. (**A**): Schematic of Wasserstein distance framework. Example quantile-quantile (QQ) plot of *p*-values from burden tests using deleterious and benign labels from CADD v1.6 to define variant masks. Signal separation measures the distance between deleterious- and benign-masked test statistics, and calibration error measures the distance between benign-masked test statistics and the null. (**B**): Calibration error and signal separation for four primary association tests using variant masks defined by one of five annotation methods, averaged across 14 quantitative traits. Points are shaped by test and colored by annota-tion.

This analysis revealed a landscape of trade-offs among techniques: annotation methods with more permissive deleterious labeling, like CADD, achieved greater signal separation (mean *W*_1_ = 14.4–15.2) than methods based on sequence models (AM, ESM, and GPN; mean *W*_1_ = 13.0–14.2) (**Fig. 3**; trait-level results in **Figs. S8, S9, S10, S11**). Among sequence models, tests using AlphaMissense had modestly higher signal separation but at the cost of poorer calibration (mean *W*_1_ = 0.07–0.15; **Fig. 3**), consistent with its higher genomic inflation in test using benign-labeled variant sets (**Fig. 2**). Among statistical tests, SKAT-O achieved the highest signal separation (mean *W*_1_ = 13.4–15.2) while burden tests maintained the lowest calibration error (mean *W*_1_ = 0.05–0.07). Three-way ANOVA confirmed that both annotation method (*F* = 5.1, 14.9, *p* = 5.0 × 10*^−^*^4^, 1.7 × 10*^−^*^11^) and statistical test (*F* = 5.6, 4.0, *p* = 9.0 × 10*^−^*^4^, 7.5 × 10*^−^*^3^) significantly affected calibration error and signal separation (**Tables S7, S10**). Pairwise comparisons with Tukey’s HSD test for both *W*_1_ metrics confirmed that tests using AlphaMissense had higher calibration error (**Table S8**) and tests using either CADD version had higher signal separation (**Table S11**), while burden tests had the lowest calibration error among association tests (**Table S9**).

### Signal validation through constraint enrichment and replication

Next, we considered the extent to which the association tests prioritized genes with relevant functional and biological signal as opposed to merely having an increased false positive rate. In the absence of a gold-standard truth set of causal genetic effects, we performed three types of validation to further interpret results from statistical association tests using each annotation method. First, we examined signal enrichment in loss-of-function (LoF) intolerant genes, which have been shown to harbor an excess of rare-variant heritability [4]. Second, we sought to replicate gene–trait associations in pairs of highly related phenotypes. Third, we conducted the same replication using burden tests based on LoF variants [26].

To examine enrichment in LoF-intolerant genes, we considered genes in the top and bottom genome-wide quartiles of *s*_het_ (computed using GeneBayes [27]), which measures selection against heterozygous loss-of-function variation. For each annotation method and statistical test, we used the set of nominally significant gene–trait associations (*P* ≤ 10*^−^*^4^) to compute an enrichment ratio as the number of constrained genes (largest quartile of *s*_het_) over the number of unconstrained genes (**Fig. 4**; **Methods**).

**Fig. 4:**
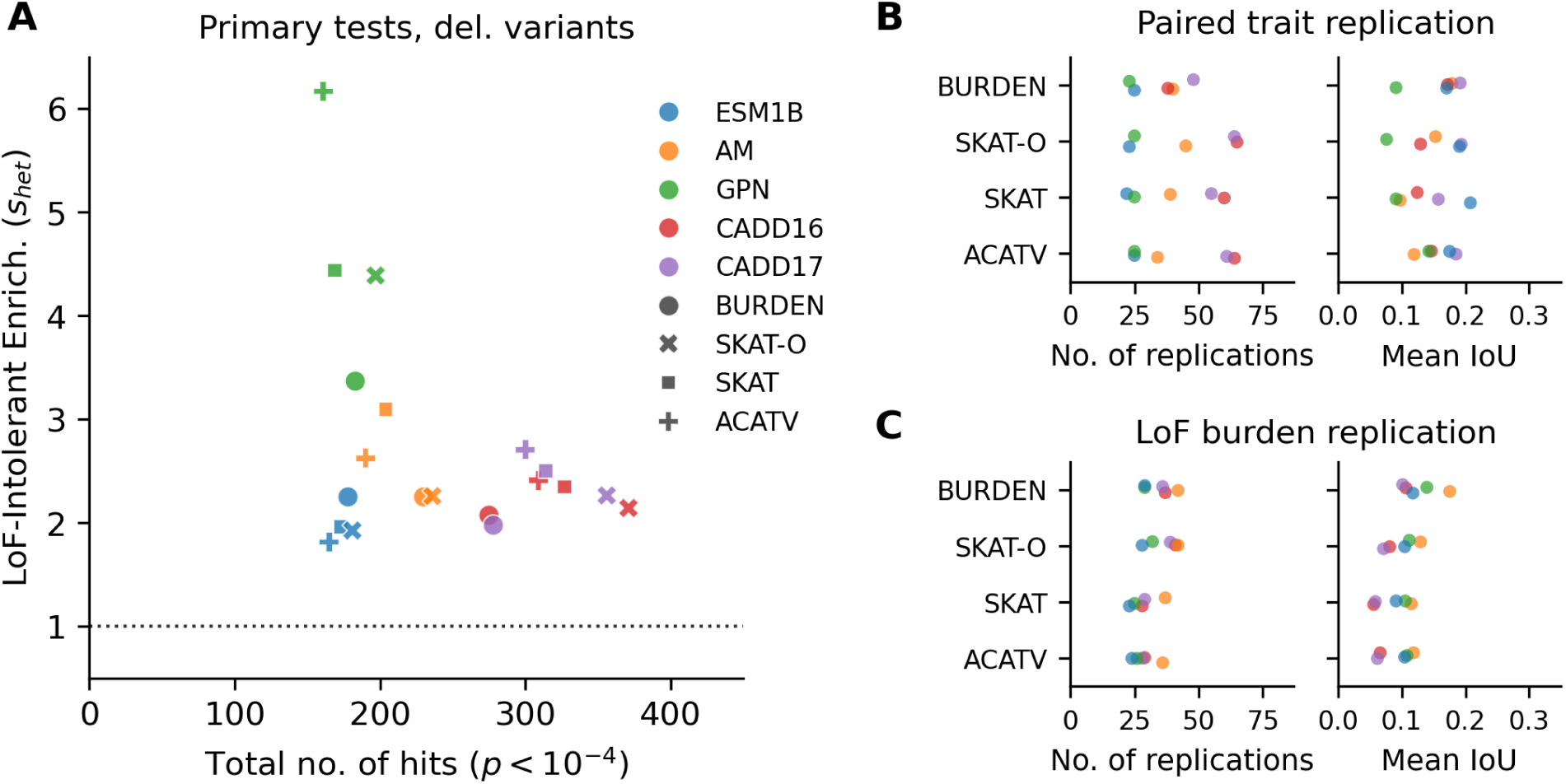
Signal validation in primary tests. For each primary test, (**A**) the total number of significant gene-trait pairs with *p <* 10*^−^*^4^ summed across all 14 traits, and the enrichment of those genes in the top quartile (25%) of *s*_het_ compared to the bottom quartile; (**B**) the total number of genes that are replicated hits (*p <* 10*^−^*^4^) and the mean intersection over union (IoU) of sets that are hits across eight closely related trait pairs; and (**C**) the total number of replicated hits and IoU scores in loss of function burden tests for five traits.

Significant results from primary tests using deleterious masks showed 1.8–5.8-fold enrichment in constrained genes across all annotation methods, compared to 0.9–3.4-fold for moderate and 0.6–1.9-fold for benign masks (**Fig. S12**). GPN-MSA achieved the highest enrichment ratios (up to 5.8-fold for ACAT-V; **Fig. 4A**, **Tables S13** and **S14**), consistent with its stringent deleteriousness classification and with its variant scores exhibiting the strongest correlation with gene-level LoF intolerance (*s*_het_ and LOEUF) among all annotation methods (**Fig. S13**). We found this elevated enrichment ratio to be consistent across *p*-value thresholds (**Fig. S14**). There was otherwise comparable enrichment of LoF-intolerant genes among significant associations across primary tests using other annotation methods, even as these approaches produced different numbers of nominally significant hits (**Fig. 4A**, **Tables S16**, **S17**, and **S18**).

Next, we sought to validate the gene–trait pairs prioritized by these methods using trait- and variant-level consistency checks. At the trait level, our evaluation set contains eight pairs of strongly correlated traits between which we expected to find meaningfully overlapping signal (**Methods**). These include left- and right-eye measurements of ocular phenotypes [IOP, IOPg, CH, and CRF], and anthropometric ratios [BMI, FEV1/FVC] where pleiotropy between the ratio and its constituent traits is expected given sufficient power [28]. Across annotation methods and statistical tests, we found that the number of genes nominated as hits for both traits across pairs varied in a manner consistent with the overall number of gene–trait associations at this significance threshold (*P <* 10*^−^*^4^; **Fig. 4B**). We also observed the same relative performance of annotation method and statistical tests using this measure, with tests using CADD annotations producing more replicated signal. However, we observed no relationship between the fraction of overlapping gene–trait associations when computing the intersection over union (IoU score; **Methods**) of associated genes across trait pairs (**Fig. 4B**).

Finally, at the variant level, we compared the overlap between genes nominated for five traits — height, weight, BMI, FEV1, and FVC — with those prioritized by gene loss-of-function burden tests [26], which were computed using a disjoint set of variants compared to our analysis. As with the paired trait analysis, we found a variable but substantial overlap (counts) in genes nominated using tests that aggregate variants with different functional consequences, and that this overlap was consistent and low as a proportion of overall signal (IoU scores; **Fig. 4C,F**). We again find that tests using CADD annotations have the highest numbers of overlapping genes, and that no annotation method or statistical test has a significantly better replication rate. Taken together, these results suggest that performance differences across annotation methods and statistical tests owe mainly to power improvements resulting from the use of permissive deleteriousness labels.

### Secondary association tests minimize effects of annotation choice

We also evaluated the results of secondary association tests which aggregate signal across benign, moderate and deleterious variant masks. We considered the same criteria as for primary tests: signal enrichment in LoF-intolerant genes, replication across trait pairs, and replication with LoF burden tests.

As with primary tests, we observed more hits in the constrained genome, where with the exception of COAST_O, hits were enriched 1.6–3.2-fold in LoF-intolerant genes (**Fig. 5A**). However, unlike for primary tests, no annotation method produced a statistically measurable change in the number of hits or the constraint enrichment of prioritized genes — these differences were related mainly to the choice of association test, which were broadly stratified by their modeling assumptions (ANOVA and Tukey HSD tests; **Tables S19–S24**). The allelic series test (COAST_O) had the fewest hits and no enrichment to LoF-intolerant genes; tests that assumed genes have consistent directions of effect (BURDEN-ACAT and BURDEN-SBAT) had more hits, which showed enrichment to constrained genes; and tests using variance components (ACATV-ACAT, SKATO-ACAT, and GENE_P) had the most hits, but these hits did not have stronger constraint enrichment than those from the burden tests (**Fig. 5A**). We find the same result when using the loss-of-function observed-expected upper-bound fraction (LOEUF) as a measure of gene constraint in place of *s*_het_ (**Fig. S15**; **Methods**).

**Fig. 5:**
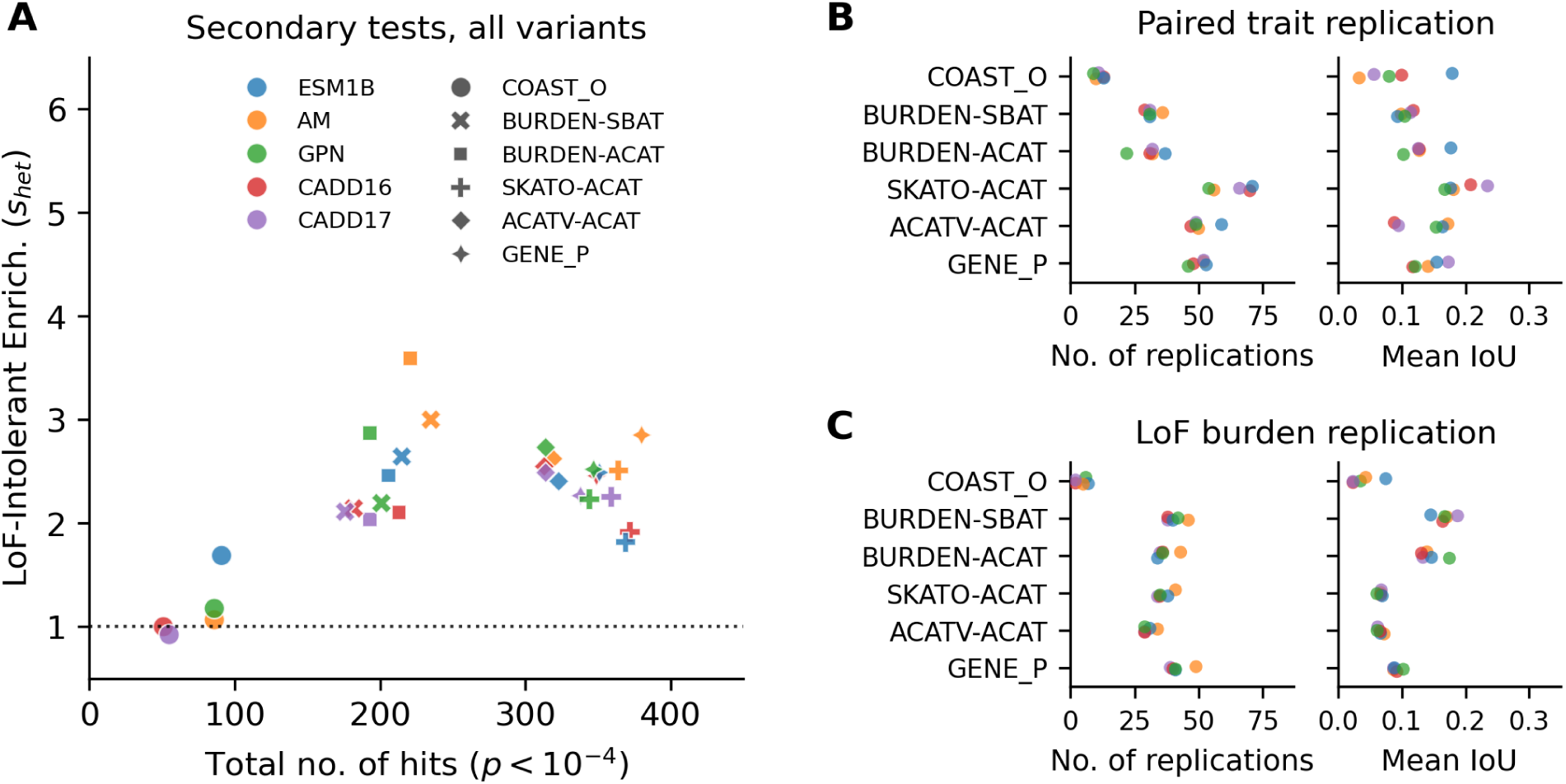
Signal validation in secondary tests. For each secondary test, (**A**) the total number of significant gene-trait pairs with *p <* 10*^−^*^4^ summed across all 14 traits, and the enrichment of those genes in the top quartile (25%) of *s*_het_ compared to the bottom quartile; (**B**) the total number of genes that are replicated hits (*p <* 10*^−^*^4^) and the mean intersection over union (IoU) of sets that are hits across eight closely related trait pairs; and (**C**) the total number of replicated hits and IoU scores in loss of function burden tests for five traits.

Across both replication sets (paired traits and LoF burden tests; **Fig. 5B** and **5C** respectively), we find the same stratification of performance by test assumption, with no statistically significant effects due to annotation method. For paired traits, COAST_O had the least replicated signal, BURDEN-ACAT and BURDEN-SBAT had a middling amount, and SKATO-ACAT, ACATV-ACAT, and GENE_P had the most replications (**Fig. 5B**). However, the IoU scores for all of these tests were broadly similar, suggesting that this trend is driven mainly by differences in power (**Fig. 5B**). In LoF burden tests, only COAST_O had a noticeably lower number of replications, and the burden-based methods had a higher replication rate (**Fig. 5C**), which may simply reflect that they make the same assumptions as tests used for comparison rather than indicate superior performance. We find no significant effect of annotation method on either statistic across both replication sets, suggesting that secondary tests collapse differences in signal that otherwise arises in analyses of subsets of the input variant set (e.g., when considering only variants labeled as deleterious, as in **Fig. 4**).

## Discussion

In this work, we systematically evaluated the use of machine-learning based variant annotation methods for rare variant association testing. Our findings reveal trade-offs between statistical power and calibration that inform methodological choices, providing actionable guidance for future studies and introducing a new framework for assessing the performance of these tools.

Annotation methods showed substantial differences in variant classification that translated into measurable performance impacts on primary association tests. Tests using the more permissive deleteriousness labels from ensemble-based annotation methods (CADD) achieved the highest power, while tests using more conservative labels from sequence models like AlphaMissense showed poorer calibration. Tests using labels from GPN-MSA produced results with the strongest enrichment in LoF constrained genes. Across primary statistical tests, burden and hybrid methods (SKAT-O) maintained the best calibration, as previously observed with point estimates of genomic inflation [29, 30], while none of the approaches was clearly the most or least powerful. Secondary tests, which aggregate signal across labels of variant deleteriousness, largely erased differences across annotation methods as all scored variants were considered, in one category or another, by these statistical tests. As in prior work [16, 19, 31], we found the power of these tests to be tied to their modeling assumptions — tests based on variance components had stronger signal than burden tests assuming a consistent direction of effect, which in turn had more signal than an allelic series test (COAST). These secondary tests also had marked differences in calibration, although they generally had similar levels of constraint enrichment.

Taken together, our findings show that no single combination of annotation method and statistical test is optimal across performance criteria. While study priorities can inform the selection of a statistical test (e.g., via the need to estimate a magnitude and direction of gene effects), differences between annotation methods are more subtle to interpret. Since these methods generally produced concordant variant scores, we suspect that resulting performance differences may be a function of the thresholds used to label variants as benign, moderate, or deleterious. As a result, we contend that this practice of binning variants deserves closer scrutiny in the context of association testing. At minimum, the boundaries used to define inclusion in association tests could be redrawn to improve on existing guidelines, which, for CADD, are rank-based [8, 9], and for sequence models, have been calibrated against benchmarks in clinical variant classification [10, 12, 14].

We also acknowledge limitations in our study design. Our analysis used data on quantitative traits from European ancestry participants in the UK Biobank, and the relative performance of method combinations may differ in association tests of binary outcomes, in other cohorts, or with tests that make different use of model-based pathogenicity scores. Our constraint enrichment analysis, while supportive of biological relevance, represents only one dimension of functional validation. Our validation experiments across related traits and with loss-of-function variants, while informative of test performance, are not comprehensive, and applications of our approach to other traits and cohorts will be insightful. Similarly, the variant masks we use for testing in this work used hard filters for allele count and frequency: these choices may also influence our results, and we encourage further investigation into the (possibly nonlinear) relationships between variants’ pathogenicity scores, allele frequencies, and effects on complex traits and diseases [6].

Finally, we note that while all of the studied annotation methods capture some aspect of variants’ effects on genes, their relationship to genes’ effects on fitness (e.g., selective constraint against heterozygous loss of function) is much more variable (**Fig. S16**). Importantly, rare-variant heritability is observed to be enriched among constrained genes [4], and the nature of a variant’s relationship to a gene, a gene’s relationship to a trait, and each of their respective fitness consequences are all key determinants of power and ranking in association studies [5]. More thoughtful use of measures of the functional consequences of variants and genes will therefore be critical for improving variant inclusion criteria in future rare-variant association studies.

## Methods

### Study population and phenotypes

We analyzed whole-exome sequencing data from the UK Biobank, using a set of up to 350,377 participants of European ancestry. We considered two analysis cohorts: one, of unrelated individuals excluding up to third-degree relatives (the “unrelated” cohort, *n* = 297,617), and another including related individuals (the “all” cohort, *n* = 350,377). We analyzed 14 quantitative traits in three categories: anthropometric (height, weight, and body mass index [BMI]), pulmonary function (forced vital capacity [FVC], forced expiratory volume in one second [FEV1], and a derived FEV1/FVC ratio), and ocular measurements (intraocular pressure with corneal compensation and Goldman cor-relation [IOP, IOPg], corneal hysteresis [CH], corneal resistance factor [CRF] for left and right eyes). All phenotypes were inverse rank normalized and all association tests were adjusted for age, sex, age^2^, age×sex, age^2^×sex, assessment center, and the first 20 principal components of the genotype matrix.

### Variant annotation

We evaluated five machine learning-based annotation methods which are commonly used for variant pathogenicity prediction and whose licenses permitted our analysis. Briefly, these include two versions of Combined Annotation-Dependent Depletion scores (CADD v1.6 [8] and v1.7 [9]), which are derived from a logistic regression model of variant-level features spanning genomic annotations and functional information; AlphaMissense (AM) [10], which is a deep-learning model of missense variant pathogenicity based on AlphaFold2 [32] and fine-tuned on human population allele frequency data; ESM-1b [11], which is a protein language model using the transformer neural network architecture trained on protein sequence data from multiple species; and GPN-MSA [12], which is a DNA language model using transformers trained on aligned genomic sequences from multiple species.

Our analyses focused on the complete set of 9,335,541 single nucleotide variants annotated as protein-coding in the gnomAD v4.1 resource [15] (this number separately counts each alternate for multi-allelic loci). Variants were scored by each of the above methods and then classified into three categories (benign, moderate, or deleterious) using method-specific thresholds previously established in the literature. For both CADD versions, benign variants scored below 10, moderate variants below 20, and deleterious variants above 20 (phred-scale) [9]; for AM, benign variants scored below 0.34, moderate variants below 0.564, and deleterious variants above 0.564 [10]; for ESM-1b and GPN, benign variants scored above -3, moderate variants above -8, and deleterious variants below -8 [12, 14]. For methods that only score missense variants (AM and ESM-1b), we applied the benign label to all synonymous variants. For variants located in the protein-coding regions of multiple genes, we used their gene-specific annotations (where available) when defining variant masks for association tests, but retained their most severe consequence (e.g., missense over synonymous) or annotation (e.g., deleterious over benign) when describing variant attributes and annotations using a single label (e.g., as in **Fig. 1**).

### Rare variant association testing

We considered four primary tests that collapse signal across variants with a particular annotation (benign, moderate, or deleterious) for a gene. These are the Burden test, the sequence kernel association test (SKAT) and its optimal combination with the Burden test (SKAT-O) [19], and the aggregated Cauchy association test for variants (ACAT-V) [17]. Briefly, the burden test is an aggregation test that assumes all variants in a gene have same effect on the trait; SKAT is a variance component test that assumes variant effects are sampled from the same zero-mean distribution; SKAT-O is an omnibus test that weights SKAT and Burden test statistics to optimize genome-wide power; and ACAT-V is an aggregation test that performs a frequency-weighted combinations of variant-level *p*-values.

We further considered six gene-level aggregation tests that combine results across annotation masks. These include applying the aggregate Cauchy association test (ACAT) to results from the Burden, ACAT-V, and SKAT-O tests as above (BURDEN-ACAT, ACAT-V-ACAT, and SKAT-O-ACAT) [17, 33, 23]; the sparse burden association test (SBAT) [21]; the coding variant allelic series test (COAST) [22]; and a unified test (GENE_P) that combines results from several of these approaches [21]. Briefly, the ACAT tests aggregate *p*-values from multiple variant masks per gene using the technique described above for ACAT-V; SBAT tests the joint null over all of the variant masks; COAST tests an (ordered) weighted average of the variant masks; and GENE_P performs ACAT across results from SKAT-O, ACAT-V, BURDEN-ACAT, and SBAT on multiple variant-level masks.

With the exception of COAST, all primary variant-level and secondary annotation-level tests were performed using the implementations provided by REGENIE v3.6 [23]. Briefly, we followed the two-step inference procedure as previously described: first, we computed a whole-genome regression model by ridge regression with leave-one-out cross-validation (`–step 1 –l1`) using LD-pruned variants on the UKB genotyping array. Second, we performed gene-based association tests on exome sequencing variants with linear regression on inverse rank normalized phenotypes with the following covariates regressed out: age, sex, age×sex, age^2^, assessment center (categorically encoded), the first 20 principal components of the genotype matrix, and leave-one-chromosome out (LOCO) polygenic predictions from the first step. We used variant labels from each annotation method (benign, moderate, and deleterious, as described above) to define masks for primary association tests, with additional filters on minimum minor allele count (5 for burden tests, with `--minMAC`; 10 for variance component tests, with `--vc-MACthr`) and maximum minor allele frequency (1%, with `--vc-maxAAF`). For SKAT-O, which has an additional hyperparameter *ρ* to weight SKAT and Burden test results, we used an 11-point grid of values: *ρ* ∈ {0.0, 0.1^2^, 0.2^2^, 0.3^2^, 0.4^2^, 0.5^2^, 0.6^2^, 0.7^2^, 0.8^2^, 0.9^2^, 1.0}. Sec-ondary tests, which perform additional rounds of signal aggregation across labeled variant sets, were performed as described in the REGENIE documentation using the appropriate flag to initiate computation [23].

### Calibration and power assessment

As an initial assessment of test calibration, we computed genomic inflation factors (*λ*_GC_) as the ratio of median observed *χ*^2^ statistics to the theoretical median (approximately 0.455). We performed this assessment separately for each combination of annotation method, variant mask, statistical test, cohort, and phenotype (totaling 5 × ((4 × 3) + 6) × 2 × 14 = 2,520 tests). We present these results in aggregate across traits in our primary analyses and visualizations.

We further quantified distributional calibration and signal separation with a novel approach using the 1-Wasserstein (“earth-mover’s distance”). For two empirical samples *x* ∼ *F* (·)*, y* ∼ *G*(·) of *n* statistical tests, this distance is computed as

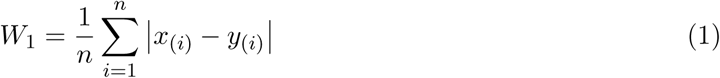

where *x*_(_*_i_*_)_ and *y*_(_*_i_*_)_ denote the *i*-th order statistics (sorted values) of each sample. This empirical estimator corresponds to the integral 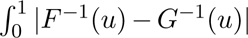 evaluated at quantile points *u* = *i/n*, and represents the average absolute difference between matched quantiles of the two distributions.

For the four primary gene-based tests, we computed this statistic on two comparison sets which serve as the basis for our assessments of calibration and power. First, we define calibration error as the *W*_1_ distance between *χ*^2^ statistics from benign-masked variants and an identically sized sample from the null 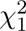 distribution. Second, we define signal separation as the *W*_1_ distance between *χ*^2^ statistics of benign- and deleterious-masked variants, restricted to tests with *p <* 10*^−^*^3^ for the deleterious mask. We did not perform this assessment for the six secondary association tests, which aggregate signal over the labels we use to define calibration error and signal separation. Instead, we evaluated these tests using point estimates like genomic inflation, and the number of nominally significant hits across traits.

### Constraint enrichment analysis

Gene constraint was quantified using estimates of selection against heterozygous loss-of-function (*s*_het_) from GeneBayes [27], and using the loss-of-function observed/expected upper bound fraction (LOEUF) from gnomAD v4.1 [34]. Genes were stratified into constrained (25th percentile) and unconstrained (75th percentile) categories for each statistic. Enrichment ratios were computed as the number of significant gene–trait associations (*P <* 10*^−^*^4^) in constrained versus unconstrained genes.

### Replication across traits and variant types

As an internal validation of the signal produced by combinations of annotation and statistical test, we assessed the extent to which the same genes were discovered in association tests for both of eight pairs of traits where we expect overlapping signal due to bilateral symmetry or the definition of a ratio trait. These trait pairs include left- and right-eye measurements of four ocular phenotypes (IOP-L, IOP-R), (IOPg-L, IOPg-R), (CH-L, CH-R), and (CRF-L, R), and four combinations of ratio traits (BMI, height), (BMI, weight), (FEV1/FVC ratio, FVC), and (FEV1/FVC ratio, FEV1). As a measure of power, we counted the number of genes that were nominally significant hits (*P <* 10*^−^*^4^) for both traits in each pair. As a measure of the rate of replication we computed the intersection over union (IoU score) for the sets of discovered genes across each trait pair.

As further validation of these gene–trait associations, we assessed signal overlap with gene loss-of-function burden tests for five traits — BMI, height, weight, FEV1, and FVC — using summary statistics from genebass [26]. We computed the same count- and intersection-based measures of overlap, described above, using these data.

## Acknowledgments

We thank Bill Forrest (Genentech) for helpful comments on the manuscript. This research has been conducted using the UK Biobank Resource under application number 44257. We thank the study participants. Aggregated summary statistics and variant scores will be made available upon publication, along with analysis code.

## Author contributions

F.J.I., M.I.M., and K.F-B. conceived the study. M.A., F.J.I., M.C., H.A.H., V.K.M., R.K.P., and K.F-B. designed analysis and/or generated data. M.A., M.C., M.I.M., and K.F-B. interpreted the results and wrote the manuscript. All authors read, revised, and approved the manuscript.

## Supplementary Tables

**Table S1:** Counts of missense and synonymous variants assigned to each category (benign, missense, and deleterious) by each annotation method (CADD = Combined annotation-dependent depletion; AM = AlphaMissense; ESM = Evolutionary scale model; GPN = Genomic Pretrained Network).

|  | Missense |  |  | Synonymous |  |  |
| --- | --- | --- | --- | --- | --- | --- |
|  | Benign | Moderate | Deleterious | Benign | Moderate | Deleterious |
| CADD16 | 910,742 | 1,346,419 | 3,961,418 | 2,394,373 | 693,047 | 29,542 |
| CADD17 | 886,162 | 1,515,204 | 3,817,213 | 2,492,336 | 599,058 | 25,568 |
| AM | 4,175,946 | 647,522 | 1,395,111 | 3,116,962 | 0 | 0 |
| ESM | 802,459 | 4,226,330 | 1,189,790 | 3,116,962 | 0 | 0 |
| GPN | 1,139,626 | 3,281,333 | 1,797,620 | 1,776,722 | 1,331,633 | 8,607 |

**Table S2:** Table of ANOVA results for three-way interactions between annotation method (AM, CADD16, CADD17, ESM, and GPN), primary association test (ACATV, BURDEN, SKAT, and SKAT-O), and study population (related and unrelated) on genomic inflation (*λ*_GC_).

| | sum_sq | df | $F$ | $P(> F)$ |
| --- | --- | --- | --- | --- |
| Annotation | 0.251 | 4 | 5.551 | $2.2 \times 10^{-4}$ |
| Test | 0.400 | 3 | 11.790 | $1.8 \times 10^{-7}$ |
| Population | 0.025 | 1 | 2.186 | $1.4 \times 10^{-1}$ |
| Annotation:Test | 0.049 | 12 | 0.361 | $9.8 \times 10^{-1}$ |
| Annotation:Population | 0.002 | 4 | 0.037 | 1.0 |
| Test:Population | 0.003 | 3 | 0.084 | $9.7 \times 10^{-1}$ |
| Annotation:Test:Population | 0.001 | 12 | 0.011 | 1.0 |

**Table S3:** Table of Tukey HSD results for multiple comparison of means (FWER=0.05) measuring effects of annotation method on genomic inflation (*λ*_GC_) in primary association tests.

| Annot. A | Annot. B | B minus A | p-adj | lower | upper | reject |
| --- | --- | --- | --- | --- | --- | --- |
| AM | CADD16 | -0.055 | 0.0013 | -0.0942 | -0.0159 | True |
| AM | CADD17 | -0.0545 | 0.0015 | -0.0936 | -0.0153 | True |
| AM | ESM | -0.0442 | 0.0179 | -0.0833 | -0.0051 | True |
| AM | GPN | -0.0542 | 0.0016 | -0.0933 | -0.015 | True |
| CADD16 | CADD17 | 0.0005 | 1.0 | -0.0386 | 0.0397 | False |
| CADD16 | ESM | 0.0108 | 0.9426 | -0.0283 | 0.05 | False |
| CADD16 | GPN | 0.0008 | 1.0 | -0.0383 | 0.04 | False |
| CADD17 | ESM | 0.0103 | 0.9521 | -0.0289 | 0.0494 | False |
| CADD17 | GPN | 0.0003 | 1.0 | -0.0388 | 0.0394 | False |
| ESM | GPN | -0.01 | 0.957 | -0.0491 | 0.0292 | False |

**Table S4:** Table of Tukey HSD results for multiple comparison of means (FWER=0.05) measuring effects of primary association test on genomic inflation (*λ*_GC_).

| Test A | Test B | B minus A | p-adj | lower | upper | reject |
| --- | --- | --- | --- | --- | --- | --- |
| ACATV | BURDEN | -0.0561 | 0.0001 | -0.0887 | -0.0236 | True |
| ACATV | SKAT | 0.0053 | 0.9749 | -0.0272 | 0.0379 | False |
| ACATV | SKAT-O | -0.0437 | 0.0033 | -0.0762 | -0.0112 | True |
| BURDEN | SKAT | 0.0615 | 0.0 | 0.0289 | 0.094 | True |
| BURDEN | SKAT-O | 0.0124 | 0.7583 | -0.0201 | 0.045 | False |
| SKAT | SKAT-O | -0.049 | 0.0007 | -0.0816 | -0.0165 | True |

**Table S5:** Table of ANOVA results for three-way interactions between annotation method (AM, CADD16, CADD17, ESM, and GPN), secondary association test (ACATV-ACAT, BURDEN-ACAT, BURDEN-SBAT, SKATO-ACAT, GENE_P, and COAST), and study population (related and unrelated) on genomic inflation (*λ*_GC_).

| | sum_sq | df | $F$ | $P(> F)$ |
| --- | --- | --- | --- | --- |
| Annotation | 0.026 | 4 | 0.245 | $9.1 \times 10^{-1}$ |
| Test | 10.994 | 5 | 82.451 | $1.6 \times 10^{-69}$ |
| Population | 0.272 | 1 | 10.216 | $1.4 \times 10^{-3}$ |
| Annotation:Test | 0.027 | 20 | 0.051 | 1.0 |
| Annotation:Population | 0.001 | 4 | 0.013 | 1.0 |
| Test:Population | 0.056 | 5 | 0.418 | $8.4 \times 10^{-1}$ |
| Annotation:Test:Population | 0.003 | 20 | 0.005 | 1.0 |

**Table S6:**
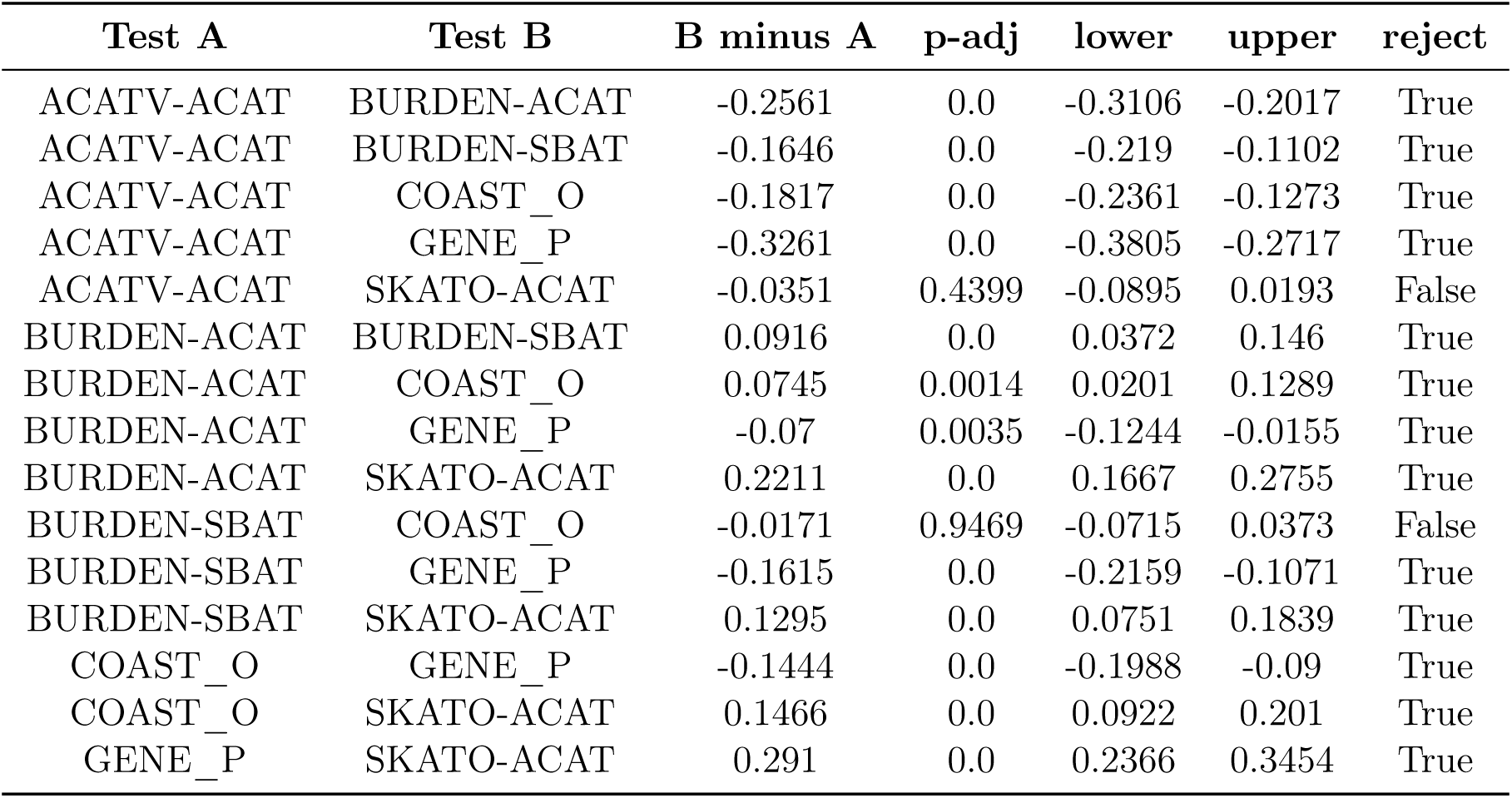
Table of Tukey HSD results for multiple comparison of means (FWER=0.05) measuring effects of secondary association test on genomic inflation (*λ*_GC_).

**Table S7:** Table of ANOVA results for three-way interactions between annotation method (AM, CADD16, CADD17, ESM, and GPN), primary association test (ACATV, BURDEN, SKAT, and SKATO), and study cohort on *W*_1_ calibration error.

| | sum_sq | df | $F$ | $P(> F)$ |
| --- | --- | --- | --- | --- |
| Annotation | 0.329 | 4 | 5.081 | $5.0 \times 10^{-4}$ |
| Test | 0.271 | 3 | 5.579 | $9.0 \times 10^{-4}$ |
| Population | 0.047 | 1 | 2.892 | $9.0 \times 10^{-2}$ |
| Annotation:Test | 0.036 | 12 | 0.186 | 1.0 |
| Annotation:Population | 0.004 | 4 | 0.056 | $9.9 \times 10^{-1}$ |
| Test:Population | 0.003 | 3 | 0.070 | $9.8 \times 10^{-1}$ |
| Annotation:Test:Population | 0.000 | 12 | 0.002 | 1.0 |

**Table S8:**
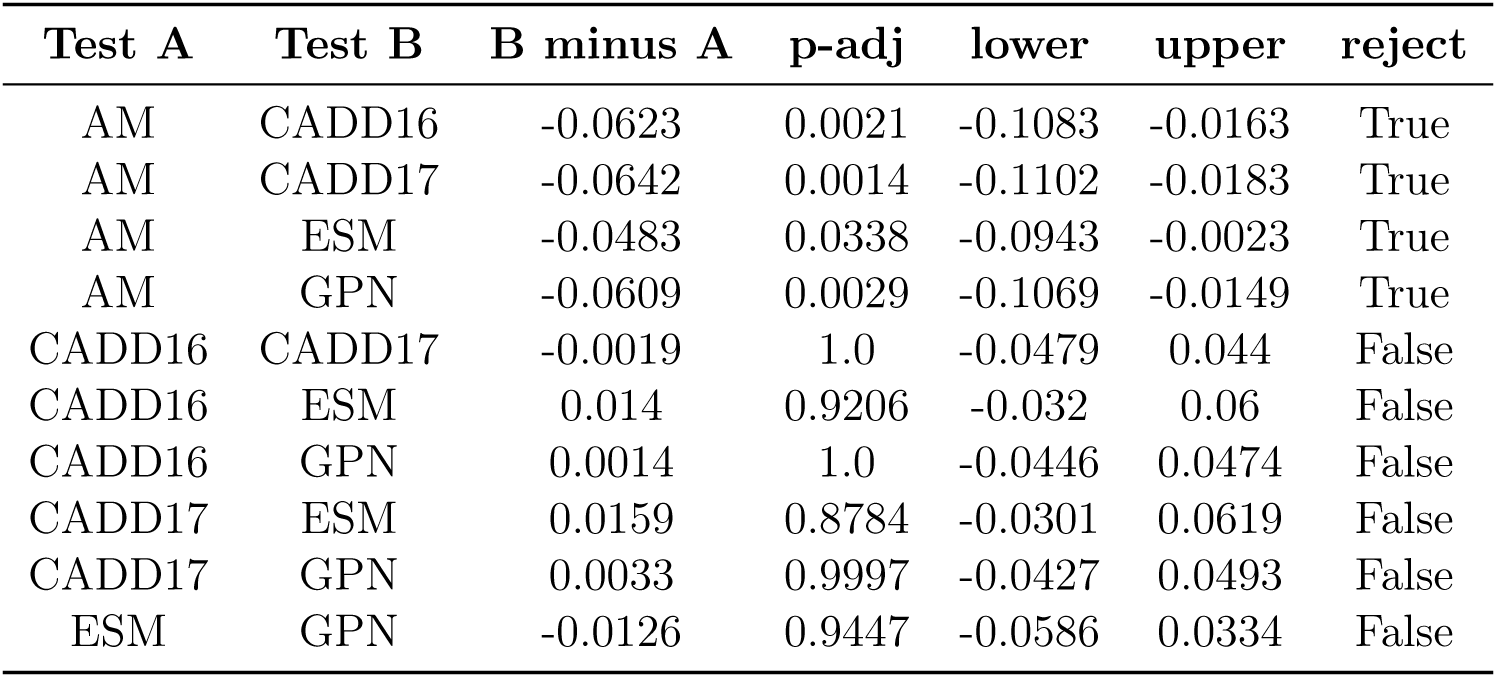
Table of Tukey HSD results for multiple comparison of means (FWER=0.05) measuring effects of annotation method on *W*_1_ calibration error.

| Test A | Test B | B minus A | p-adj | lower | upper | reject |
| --- | --- | --- | --- | --- | --- | --- |
| AM | CADD16 | -0.0623 | 0.0021 | -0.1083 | -0.0163 | True |
| AM | CADD17 | -0.0642 | 0.0014 | -0.1102 | -0.0183 | True |
| AM | ESM | -0.0483 | 0.0338 | -0.0943 | -0.0023 | True |
| AM | GPN | -0.0609 | 0.0029 | -0.1069 | -0.0149 | True |
| CADD16 | CADD17 | -0.0019 | 1.0 | -0.0479 | 0.044 | False |
| CADD16 | ESM | 0.014 | 0.9206 | -0.032 | 0.06 | False |
| CADD16 | GPN | 0.0014 | 1.0 | -0.0446 | 0.0474 | False |
| CADD17 | ESM | 0.0159 | 0.8784 | -0.0301 | 0.0619 | False |
| CADD17 | GPN | 0.0033 | 0.9997 | -0.0427 | 0.0493 | False |
| ESM | GPN | -0.0126 | 0.9447 | -0.0586 | 0.0334 | False |

**Table S9:** Table of Tukey HSD results for multiple comparison of means (FWER=0.05) measuring effects of primary association test on *W*_1_ calibration error.

| Test A | Test B | B minus A | p-adj | lower | upper | reject |
| --- | --- | --- | --- | --- | --- | --- |
| ACATV | BURDEN | -0.0503 | 0.005 | -0.0891 | -0.0115 | True |
| ACATV | SKAT | 0.0018 | 0.9994 | -0.037 | 0.0406 | False |
| ACATV | SKAT-O | -0.0004 | 1.0 | -0.0392 | 0.0384 | False |
| BURDEN | SKAT | 0.0521 | 0.0033 | 0.0133 | 0.0909 | True |
| BURDEN | SKAT-O | 0.0499 | 0.0055 | 0.0111 | 0.0887 | True |
| SKAT | SKAT-O | -0.0022 | 0.9989 | -0.041 | 0.0366 | False |

**Table S10:**
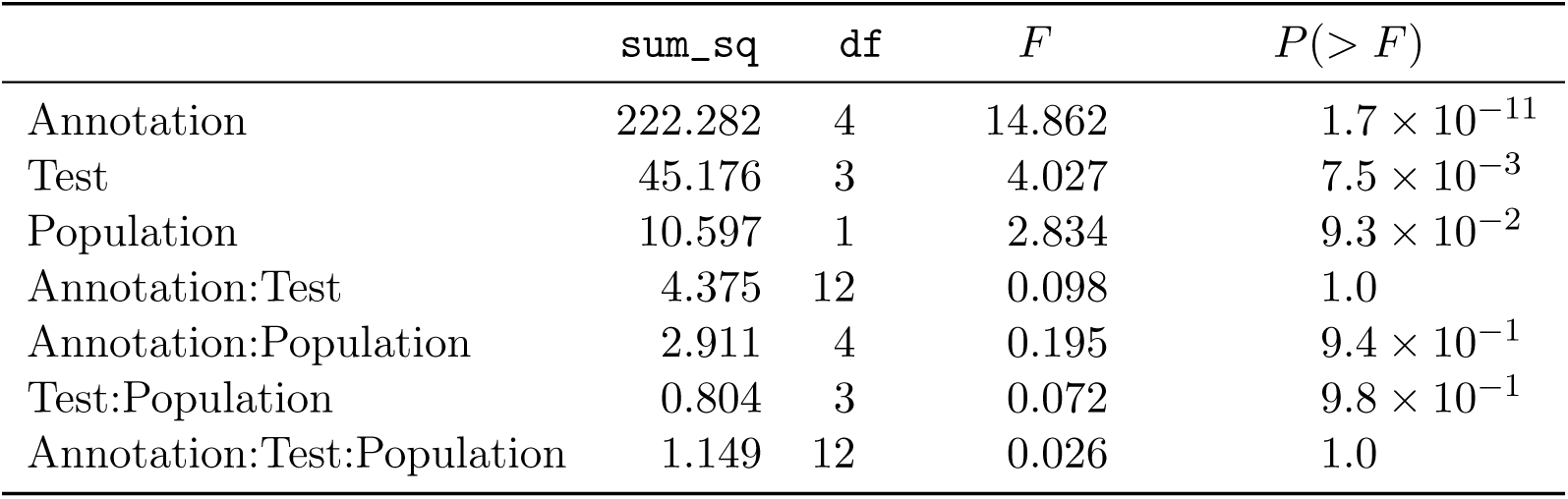
Table of ANOVA results for three-way interactions between annotation method (AM, CADD16, CADD17, ESM, and GPN), primary association test (ACATV, BURDEN, SKAT, and SKATO), and study cohort on *W*_1_ signal separation.

**Table S11:** Table of Tukey HSD results for multiple comparison of means (FWER=0.05) measuring effects of annotation method on *W*_1_ signal separation.

| Test A | Test B | B minus A | p-adj | lower | upper | reject |
| --- | --- | --- | --- | --- | --- | --- |
| AM | CADD16 | 1.1322 | 0.0001 | 0.4363 | 1.8281 | True |
| AM | CADD17 | 1.1677 | 0.0001 | 0.4719 | 1.8636 | True |
| AM | ESM | -0.1883 | 0.9468 | -0.8842 | 0.5075 | False |
| AM | GPN | -0.1953 | 0.9396 | -0.8912 | 0.5005 | False |
| CADD16 | CADD17 | 0.0356 | 0.9999 | -0.6603 | 0.7314 | False |
| CADD16 | ESM | -1.3205 | 0.0 | -2.0164 | -0.6246 | True |
| CADD16 | GPN | -1.3275 | 0.0 | -2.0234 | -0.6316 | True |
| CADD17 | ESM | -1.3561 | 0.0 | -2.0519 | -0.6602 | True |
| CADD17 | GPN | -1.3631 | 0.0 | -2.0589 | -0.6672 | True |
| ESM | GPN | -0.007 | 1.0 | -0.7029 | 0.6889 | False |

**Table S12:**
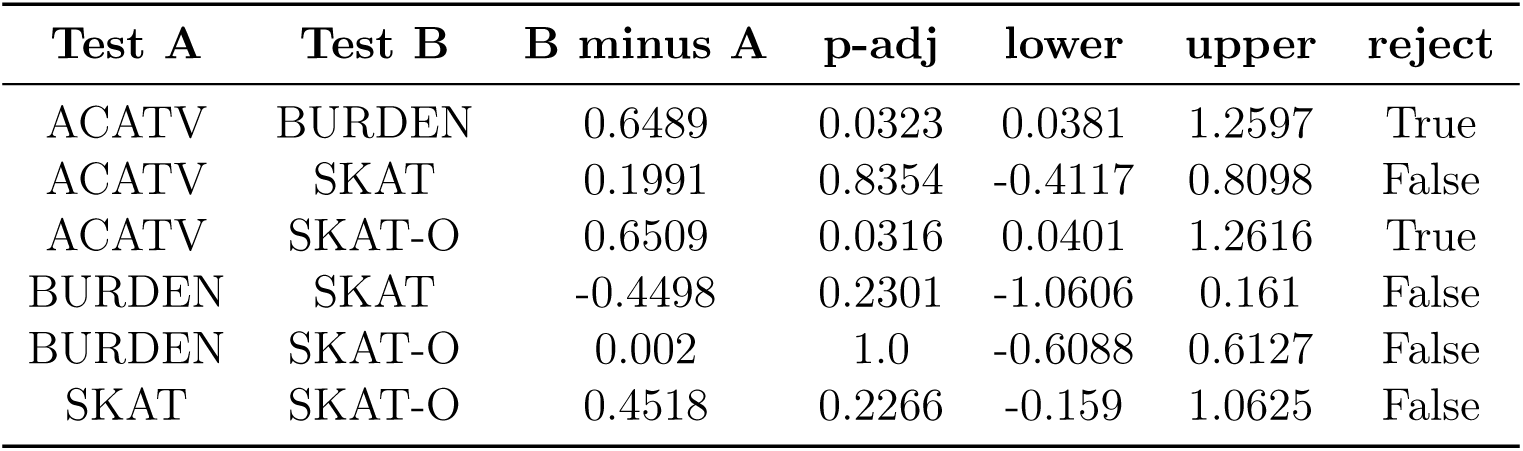
Table of Tukey HSD results for multiple comparison of means (FWER=0.05) measuring effects of primary association test on *W*_1_ signal separation.

**Table S13:** Table of ANOVA results for effects of annotation method (AM, CADD16, CADD17, ESM, and GPN) and primary association test (ACATV, BURDEN, SKAT, and SKATO) on signal enrichment in LoF-intolerant genes (top quartile of *s*_het_).

| | sum_sq | df | $F$ | $P(> F)$ |
| --- | --- | --- | --- | --- |
| Annotation | 15.807 | 4 | 14.262 | $1.6 \times 10^{-4}$ |
| Test | 1.493 | 3 | 1.796 | $2.0 \times 10^{-1}$ |

**Table S14:** Table of Tukey HSD results for multiple comparison of means (FWER=0.05) measuring effects of annotation method on signal enrichment in LoF-intolerant genes (top quartile of *s*_het_).

| Test A | Test B | B minus A | p-adj | lower | upper | reject |
| --- | --- | --- | --- | --- | --- | --- |
| AM | CADD16 | -0.331 | 0.9184 | -1.5684 | 0.9065 | False |
| AM | CADD17 | -0.0894 | 0.9994 | -1.3269 | 1.148 | False |
| AM | ESM | -0.3374 | 0.9132 | -1.5749 | 0.9 | False |
| AM | GPN | 2.0082 | 0.0012 | 0.7708 | 3.2457 | True |
| CADD16 | CADD17 | 0.2415 | 0.9725 | -0.9959 | 1.479 | False |
| CADD16 | ESM | -0.0064 | 1.0 | -1.2439 | 1.231 | False |
| CADD16 | GPN | 2.3392 | 0.0003 | 1.1018 | 3.5767 | True |
| CADD17 | ESM | -0.248 | 0.9698 | -1.4854 | 0.9895 | False |
| CADD17 | GPN | 2.0977 | 0.0008 | 0.8602 | 3.3351 | True |
| ESM | GPN | 2.3457 | 0.0003 | 1.1082 | 3.5831 | True |

**Table S15:** Table of Tukey HSD results for multiple comparison of means (FWER=0.05) measuring effects of primary association test on signal enrichment in LoF-intolerant genes (top quartile of *s*_het_).

| Test A | Test B | B minus A | p-adj | lower | upper | reject |
| --- | --- | --- | --- | --- | --- | --- |
| ACATV | BURDEN | -0.7357 | 0.7156 | -2.7143 | 1.243 | False |
| ACATV | SKAT | -0.2698 | 0.9791 | -2.2485 | 1.7088 | False |
| ACATV | SKAT-O | -0.5047 | 0.8837 | -2.4833 | 1.474 | False |
| BURDEN | SKAT | 0.4659 | 0.9055 | -1.5128 | 2.4445 | False |
| BURDEN | SKAT-O | 0.231 | 0.9867 | -1.7476 | 2.2097 | False |
| SKAT | SKAT-O | -0.2349 | 0.986 | -2.2135 | 1.7438 | False |

**Table S16:**
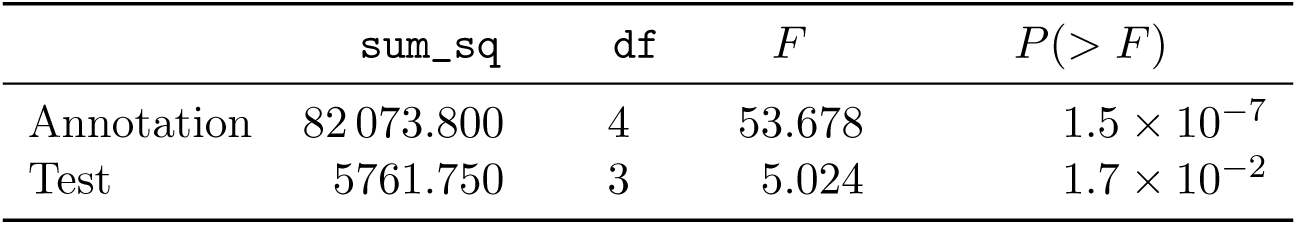
Table of ANOVA results for effects of annotation method (AM, CADD16, CADD17, ESM, and GPN) and primary association test (ACATV, BURDEN, SKAT, and SKATO) on the number of nominally significant gene-trait pairs (*p <* 10*^−^*^4^).

| | sum_sq | df | $F$ | $P(> F)$ |
| --- | --- | --- | --- | --- |
| Annotation | 82 073.800 | 4 | 53.678 | $1.5 \times 10^{-7}$ |
| Test | 5761.750 | 3 | 5.024 | $1.7 \times 10^{-2}$ |

**Table S17:** Table of Tukey HSD results for multiple comparison of means (FWER=0.05) measuring effects of annotation method on the number of nominally significant gene-trait pairs (*p <* 10*^−^*^4^) among primary association tests.

| Test A | Test B | B minus A | p-adj | lower | upper | reject |
| --- | --- | --- | --- | --- | --- | --- |
| AM | CADD16 | 105.5 | 0.0004 | 48.1478 | 162.8522 | True |
| AM | CADD17 | 97.0 | 0.0008 | 39.6478 | 154.3522 | True |
| AM | ESM | -40.75 | 0.2342 | -98.1022 | 16.6022 | False |
| AM | GPN | -37.5 | 0.3038 | -94.8522 | 19.8522 | False |
| CADD16 | CADD17 | -8.5 | 0.99 | -65.8522 | 48.8522 | False |
| CADD16 | ESM | -146.25 | 0.0 | -203.6022 | -88.8978 | True |
| CADD16 | GPN | -143.0 | 0.0 | -200.3522 | -85.6478 | True |
| CADD17 | ESM | -137.75 | 0.0 | -195.1022 | -80.3978 | True |
| CADD17 | GPN | -134.5 | 0.0 | -191.8522 | -77.1478 | True |
| ESM | GPN | 3.25 | 0.9998 | -54.1022 | 60.6022 | False |

**Table S18:**
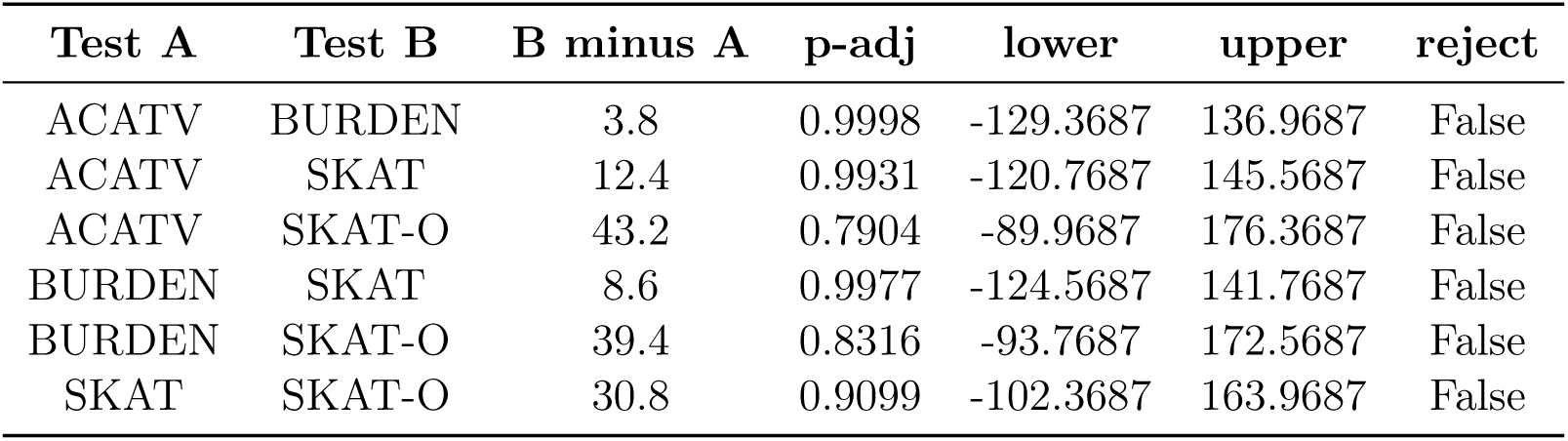
Table of Tukey HSD results for multiple comparison of means (FWER=0.05) measuring effects of primary association test on the number of nominally significant gene-trait pairs (*p <* 10*^−^*^4^).

| Test A | Test B | B minus A | p-adj | lower | upper | reject |
| --- | --- | --- | --- | --- | --- | --- |
| ACATV | BURDEN | 3.8 | 0.9998 | -129.3687 | 136.9687 | False |
| ACATV | SKAT | 12.4 | 0.9931 | -120.7687 | 145.5687 | False |
| ACATV | SKAT-O | 43.2 | 0.7904 | -89.9687 | 176.3687 | False |
| BURDEN | SKAT | 8.6 | 0.9977 | -124.5687 | 141.7687 | False |
| BURDEN | SKAT-O | 39.4 | 0.8316 | -93.7687 | 172.5687 | False |
| SKAT | SKAT-O | 30.8 | 0.9099 | -102.3687 | 163.9687 | False |

**Table S19:** Table of ANOVA results for effects of annotation method (AM, CADD16, CADD17, ESM, and GPN) and secondary association test on signal enrichment in LoF-intolerant genes (top quartile of *s*_het_).

| | sum_sq | df | $F$ | $P(> F)$ |
| --- | --- | --- | --- | --- |
| Annotation | 1.391 | 4 | 4.445 | $9.9 \times 10^{-3}$ |
| Test | 5.359 | 5 | 13.699 | $7.0 \times 10^{-6}$ |

**Table S20:** Table of Tukey HSD results for multiple comparison of means (FWER=0.05) measuring effects of annotation method on signal enrichment in LoF-intolerant genes (top quartile of *s*_het_) among secondary association tests.

| Test A | Test B | B minus A | p-adj | lower | upper | reject |
| --- | --- | --- | --- | --- | --- | --- |
| AM | CADD16 | -0.5638 | 0.3661 | -1.4561 | 0.3286 | False |
| AM | CADD17 | -0.59 | 0.3226 | -1.4823 | 0.3024 | False |
| AM | ESM | -0.2716 | 0.8964 | -1.164 | 0.6207 | False |
| AM | GPN | -0.3445 | 0.7873 | -1.2368 | 0.5478 | False |
| CADD16 | CADD17 | -0.0262 | 1.0 | -0.9185 | 0.8661 | False |
| CADD16 | ESM | 0.2921 | 0.8696 | -0.6002 | 1.1845 | False |
| CADD16 | GPN | 0.2193 | 0.9495 | -0.6731 | 1.1116 | False |
| CADD17 | ESM | 0.3183 | 0.8307 | -0.574 | 1.2107 | False |
| CADD17 | GPN | 0.2455 | 0.9258 | -0.6469 | 1.1378 | False |
| ESM | GPN | -0.0729 | 0.9992 | -0.9652 | 0.8195 | False |

**Table S21:** Table of Tukey HSD results for multiple comparison of means (FWER=0.05) measuring effects of secondary association test on signal enrichment in LoF-intolerant genes (top quartile of *s*_het_).

| Test A | Test B | B minus A | p-adj | lower | upper | reject |
| --- | --- | --- | --- | --- | --- | --- |
| ACATV-ACAT | BURDEN-ACAT | -0.0702 | 0.9995 | -0.7565 | 0.6161 | False |
| ACATV-ACAT | BURDEN-SBAT | -0.1587 | 0.9782 | -0.845 | 0.5276 | False |
| ACATV-ACAT | COAST_O | -1.2259 | 0.0001 | -1.9122 | -0.5396 | True |
| ACATV-ACAT | GENE_P | -0.1113 | 0.9956 | -0.7976 | 0.575 | False |
| ACATV-ACAT | SKATO-ACAT | -0.4559 | 0.3434 | -1.1422 | 0.2304 | False |
| BURDEN-ACAT | BURDEN-SBAT | -0.0885 | 0.9985 | -0.7748 | 0.5978 | False |
| BURDEN-ACAT | COAST_O | -1.1557 | 0.0003 | -1.842 | -0.4695 | True |
| BURDEN-ACAT | GENE_P | -0.0411 | 1.0 | -0.7274 | 0.6452 | False |
| BURDEN-ACAT | SKATO-ACAT | -0.3857 | 0.5218 | -1.072 | 0.3006 | False |
| BURDEN-SBAT | COAST_O | -1.0672 | 0.0009 | -1.7535 | -0.3809 | True |
| BURDEN-SBAT | GENE_P | 0.0474 | 0.9999 | -0.6389 | 0.7337 | False |
| BURDEN-SBAT | SKATO-ACAT | -0.2972 | 0.7611 | -0.9835 | 0.3891 | False |
| COAST_O | GENE_P | 1.1146 | 0.0005 | 0.4283 | 1.8009 | True |
| COAST_O | SKATO-ACAT | 0.7701 | 0.0216 | 0.0838 | 1.4563 | True |
| GENE_P | SKATO-ACAT | -0.3446 | 0.6356 | -1.0309 | 0.3417 | False |

**Table S22:**
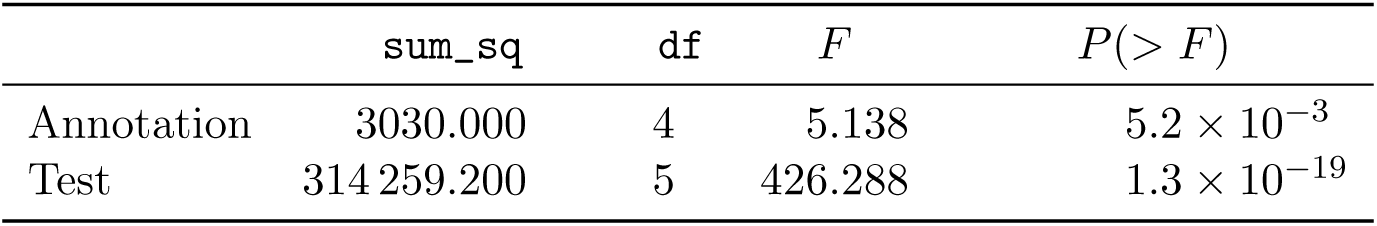
Table of ANOVA results for two-way interactions between annotation method (AM, CADD16, CADD17, ESM, and GPN) and secondary association test on the number of nominally significant gene-trait pairs (*p <* 10*^−^*^4^).

**Table S23:** Table of Tukey HSD results for multiple comparison of means (FWER=0.05) measuring effects of annotation method on the number of nominally significant gene-trait pairs (*p <* 10*^−^*^4^) among secondary association tests.

| Test A | Test B | B minus A | p-adj | lower | upper | reject |
| --- | --- | --- | --- | --- | --- | --- |
| AM | CADD16 | -20.8333 | 0.9976 | -211.8303 | 170.1636 | False |
| AM | CADD17 | -28.3333 | 0.992 | -219.3303 | 162.6636 | False |
| AM | ESM | -8.3333 | 0.9999 | -199.3303 | 182.6636 | False |
| AM | GPN | -20.0 | 0.9979 | -210.9969 | 170.9969 | False |
| CADD16 | CADD17 | -7.5 | 1.0 | -198.4969 | 183.4969 | False |
| CADD16 | ESM | 12.5 | 0.9997 | -178.4969 | 203.4969 | False |
| CADD16 | GPN | 0.8333 | 1.0 | -190.1636 | 191.8303 | False |
| CADD17 | ESM | 20.0 | 0.9979 | -170.9969 | 210.9969 | False |
| CADD17 | GPN | 8.3333 | 0.9999 | -182.6636 | 199.3303 | False |
| ESM | GPN | -11.6667 | 0.9998 | -202.6636 | 179.3303 | False |

**Table S24:** Table of Tukey HSD results for multiple comparison of means (FWER=0.05) measuring effects of secondary association test on the number of nominally significant gene-trait pairs (*p <* 10*^−^*^4^).

| Test A | Test B | B minus A | p-adj | lower | upper | reject |
| --- | --- | --- | --- | --- | --- | --- |
| ACATV-ACAT | BURDEN-ACAT | -111.4 | 0.0 | -142.2646 | -80.5354 | True |
| ACATV-ACAT | BURDEN-SBAT | -114.8 | 0.0 | -145.6646 | -83.9354 | True |
| ACATV-ACAT | COAST_O | -242.8 | 0.0 | -273.6646 | -211.9354 | True |
| ACATV-ACAT | GENE_P | 36.4 | 0.0143 | 5.5354 | 67.2646 | True |
| ACATV-ACAT | SKATO-ACAT | 45.0 | 0.0018 | 14.1354 | 75.8646 | True |
| BURDEN-ACAT | BURDEN-SBAT | -3.4 | 0.9993 | -34.2646 | 27.4646 | False |
| BURDEN-ACAT | COAST_O | -131.4 | 0.0 | -162.2646 | -100.5354 | True |
| BURDEN-ACAT | GENE_P | 147.8 | 0.0 | 116.9354 | 178.6646 | True |
| BURDEN-ACAT | SKATO-ACAT | 156.4 | 0.0 | 125.5354 | 187.2646 | True |
| BURDEN-SBAT | COAST_O | -128.0 | 0.0 | -158.8646 | -97.1354 | True |
| BURDEN-SBAT | GENE_P | 151.2 | 0.0 | 120.3354 | 182.0646 | True |
| BURDEN-SBAT | SKATO-ACAT | 159.8 | 0.0 | 128.9354 | 190.6646 | True |
| COAST_O | GENE_P | 279.2 | 0.0 | 248.3354 | 310.0646 | True |
| COAST_O | SKATO-ACAT | 287.8 | 0.0 | 256.9354 | 318.6646 | True |
| GENE_P | SKATO-ACAT | 8.6 | 0.952 | -22.2646 | 39.4646 | False |

## Supplementary Figures

**Fig. S1:**
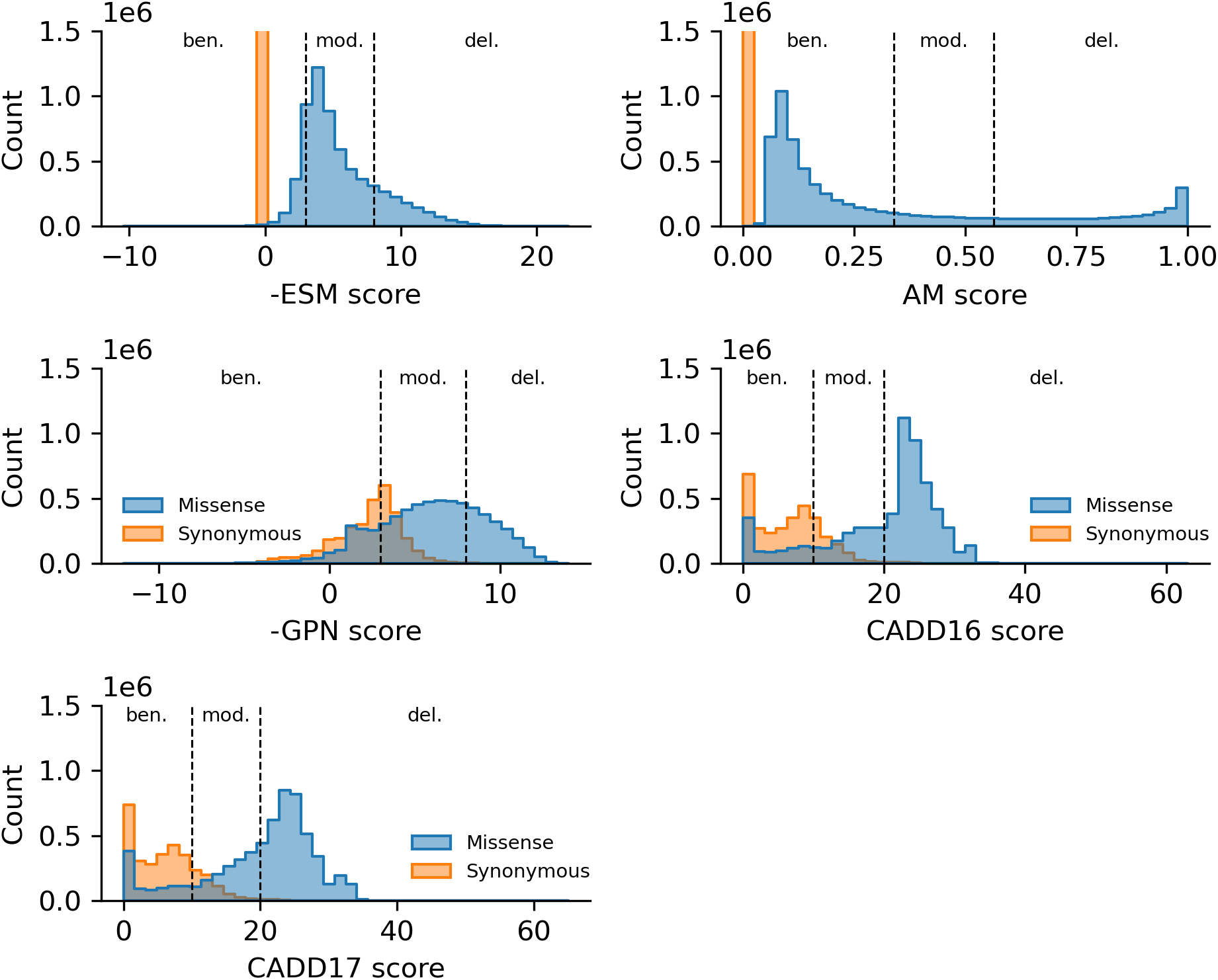
Distribution of variant scores across annotation methods and variant types. Histogram of variant scores across for annotation method (panels), stratified by variant type (missense or synonymous). In all panels, the *y*-axis is capped at 1.5 million, which truncates bars for synonymous variants in cases where we imputed zero values for these variants (AM and ESM1b only score missense variants). Scores for ESM1b and GPN are negated so as to polarize all scores with the same direction of interpretation (larger is more delterious). Each panel is also annotated with the method-specific thresholds used to define variants as benign, moderate, or pathogenic.

**Fig. S2:**
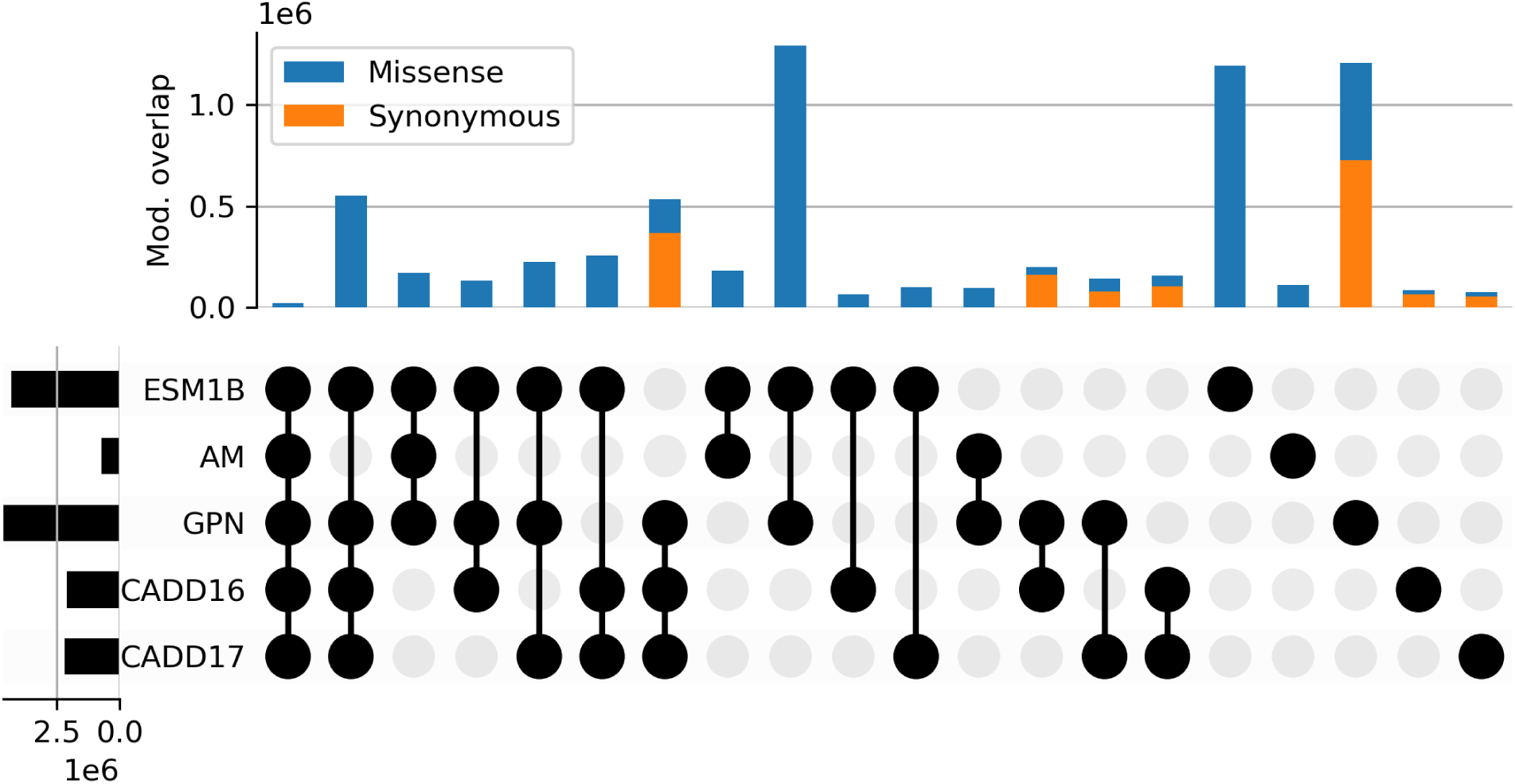
UpSet plot of overlap in variants labeled as moderate by each of the annotation methods. There are 1,550,046 synonymous and 5,310,920 missense variants labeled as moderate by any annotation method in this plot. Bars are not shown if they account for less than 0.2% of the total variant set.

**Fig. S3:**
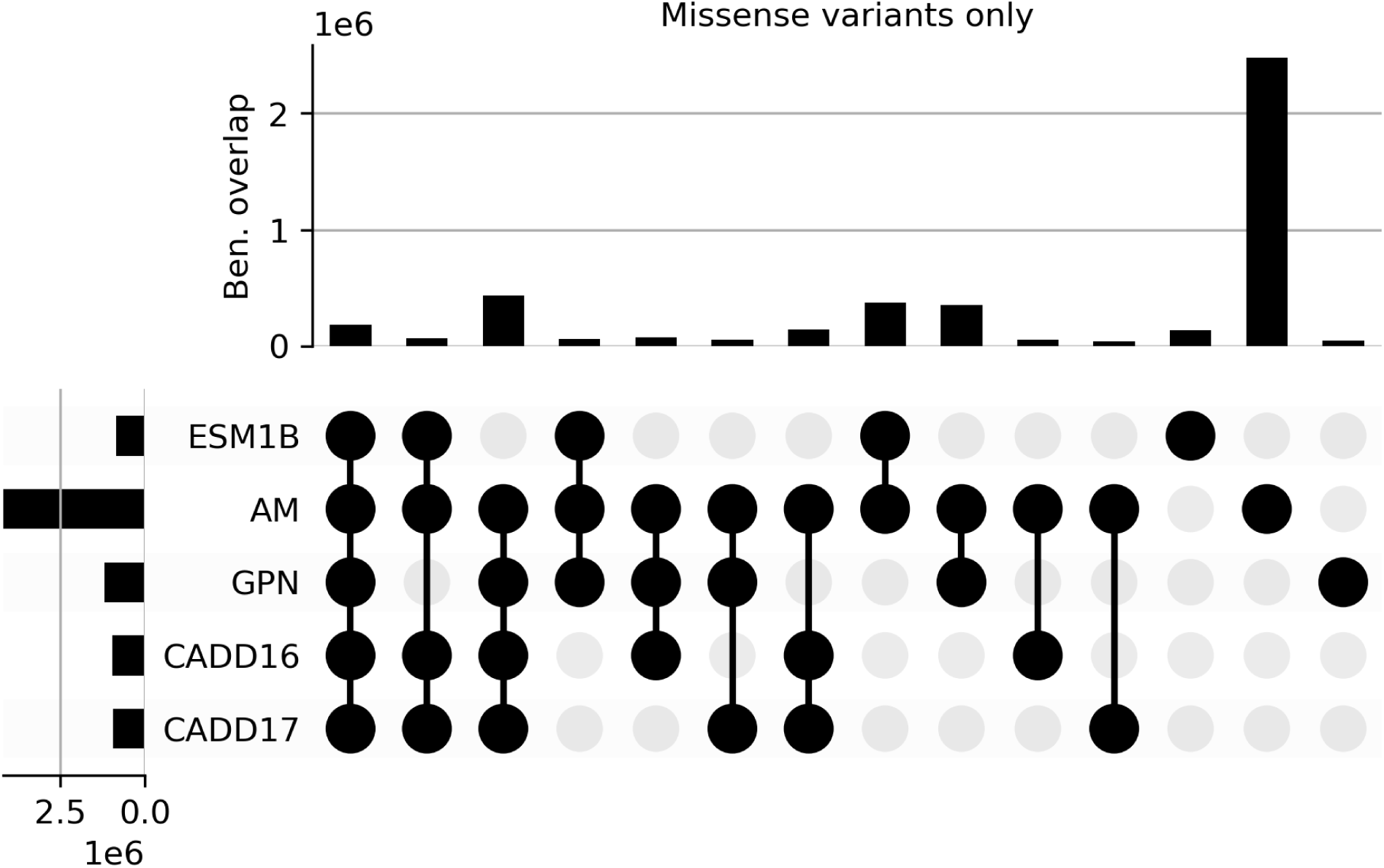
UpSet plot of overlap in missense variants labeled as benign by each of the annotation methods. There are 4,367,688 missense variants labeled as benign by any annotation method in this plot.

**Fig. S4:**
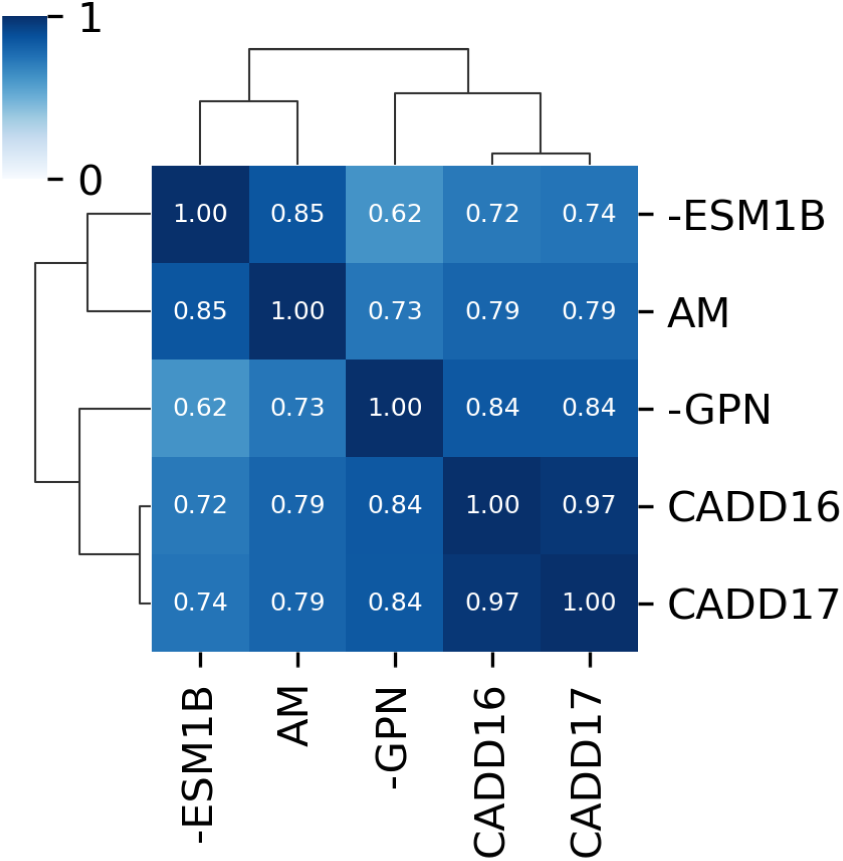
Clustered heatmap of raw deleteriousness scores from each annotation methods for all variants (synonymous variants are set to zero for ESM1B and AM; see **Methods**). Numbers in each box indicate the rank correlation (Spearman *ρ*) between pairs of methods. Scores from ESM and GPN are negated to polarize scores from all methods to the same direction (larger is more deleterious).

**Fig. S5:**
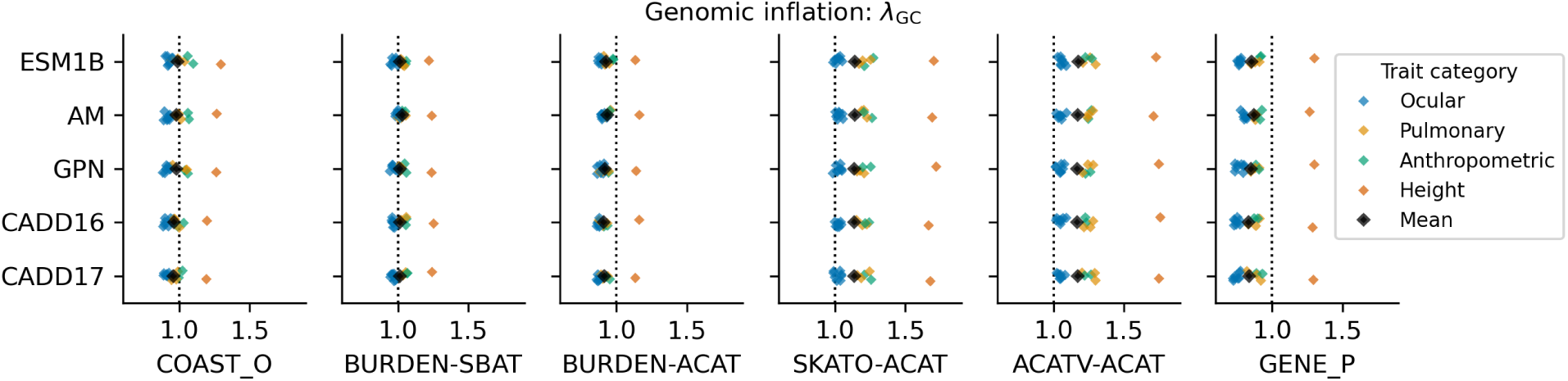
Genomic inflation by secondary test and annotation method. Genomic inflation (*λ*_GC_) of secondary test methods (subpanels) using variant classification sets produced by each of the annotation methods (*y*-axis bins). Each point corresponds to one of 14 traits × 5 annotations × 6 secondary tests = 420 RVATs, and is colored by phenotype category.

**Fig. S6:**
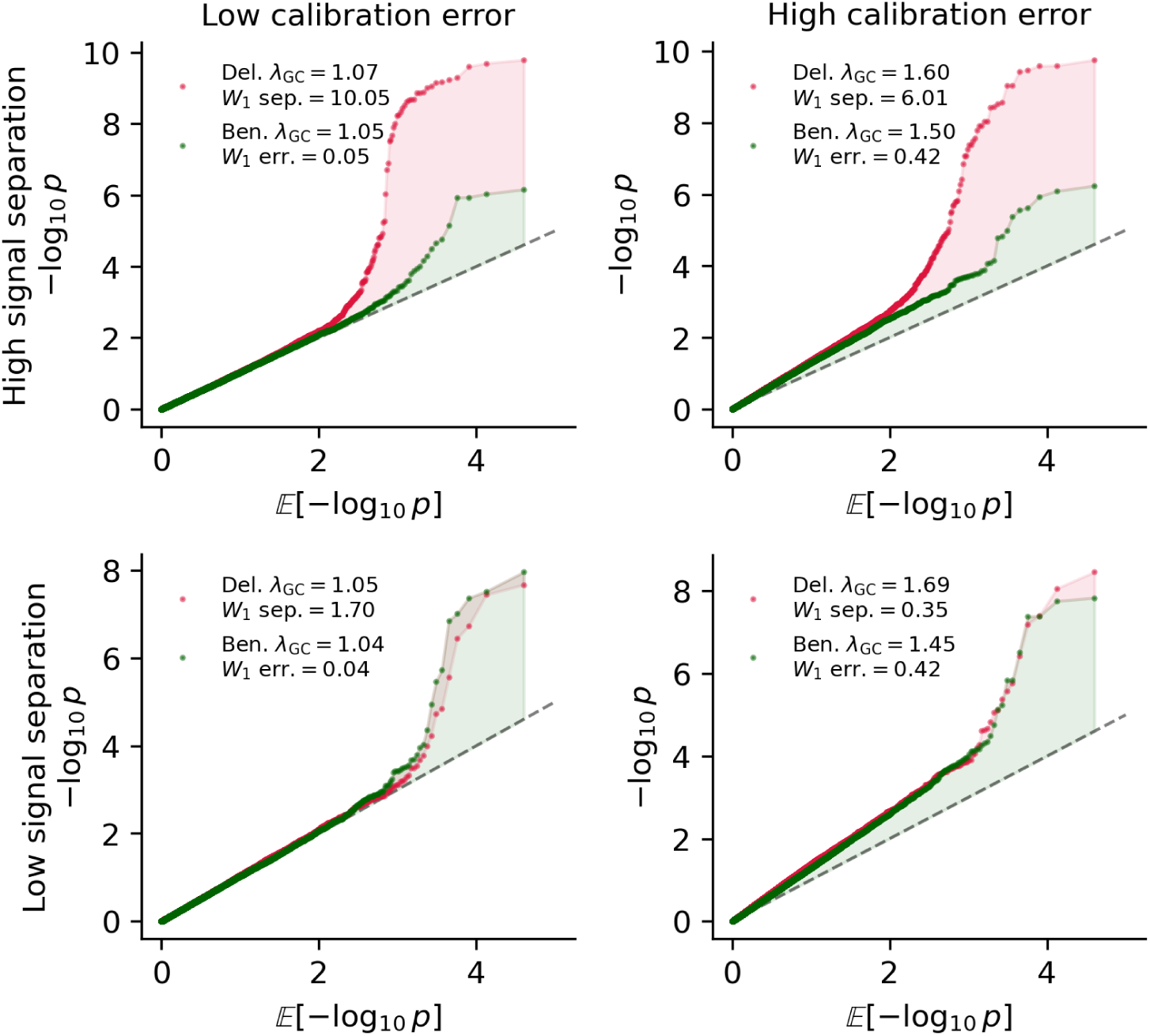
Examples of calibration error and signal separation compared to genomic inflation. Cartoon quantile-quantile plots of four synthetic association test scenarios: high or low calibration error (similar to benign genomic inflation), and high or low signal separation (similar to number of hits in the deleterious category). Each panel shows the computed genomic inflation factor for the sets of benign- and deleterious-labeled association test statistics, and the computed Wasserstein distances between the distributions (see **Methods** for formulas).

**Fig. S7:**
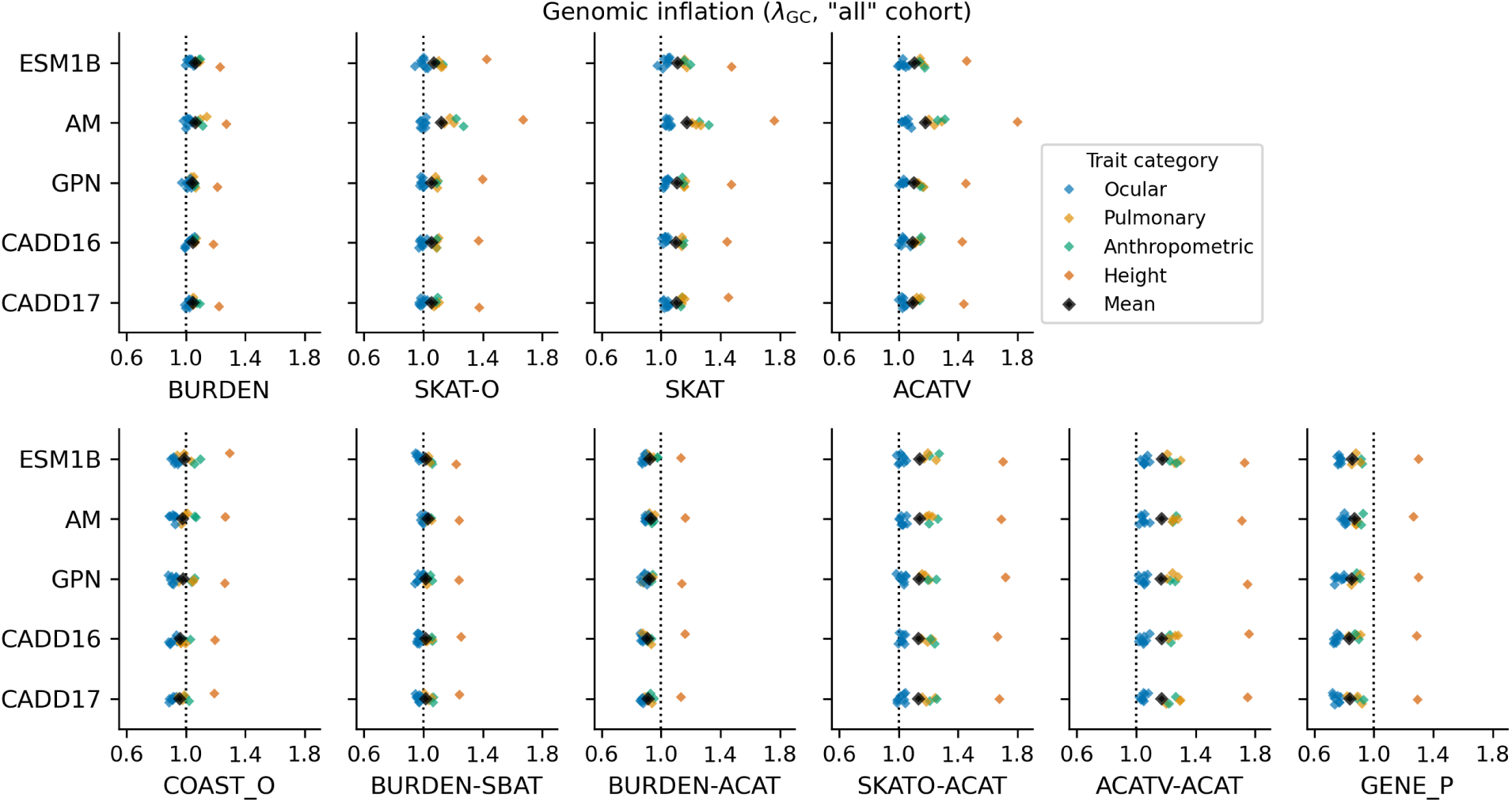
Genomic inflation by test and annotation method in the “all” cohort. Genomic inflation (*λ*_GC_) of secondary test methods (subpanels) using benign (primary, top row) or all variant classification sets produced by each of the annotation methods (*y*-axis bins). Each point corresponds to one of 14 traits × 5 annotations × 10 tests = 700 RVATs, and is colored by phenotype category.

**Fig. S8:**
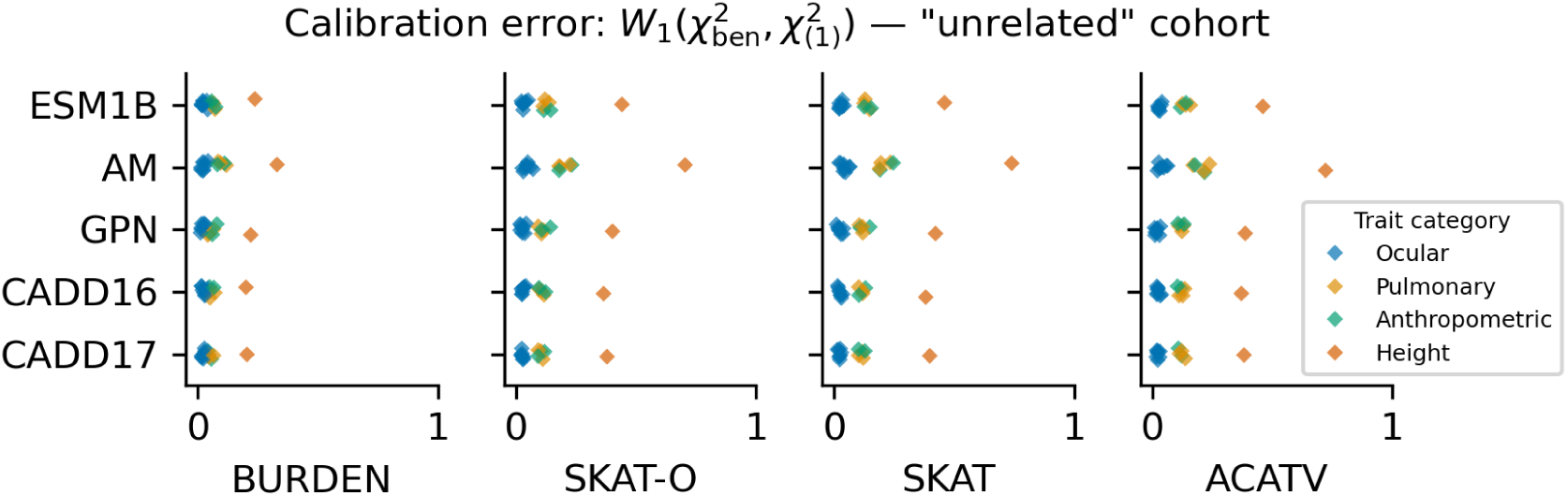
Null calibration by primary test and annotation method. Null calibration (*W*_1_ distance between benign-mask chi-squared statistics and the null 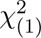 distribution) of primary test methods (subpanels) in the “unrelated” cohort using variants classified as benign by each of the annotation methods (*y*-axis bins). Each point corresponds to one of 14 traits × 5 annotations × 4 tests = 280 RVATs, and is colored by phenotype category.

**Fig. S9:**
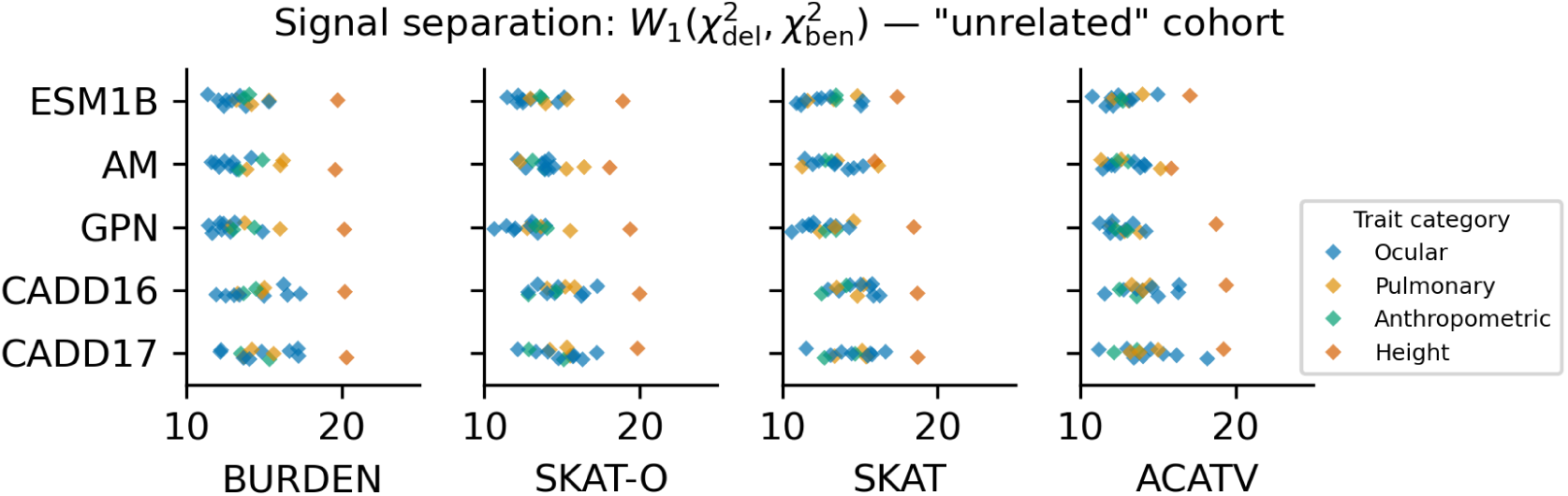
Signal separation by primary test and annotation method. Signal separation (*W*_1_ distance between benign-mask and deleterious-mask chi-squared statistics) of primary test methods (subpanels) in the “unrelated” cohort using labels from each of the annotation methods (*y*-axis bins). Each point corresponds to one of 14 traits × 5 annotations × 4 tests = 280 RVATs, and is colored by phenotype category.

**Fig. S10:**
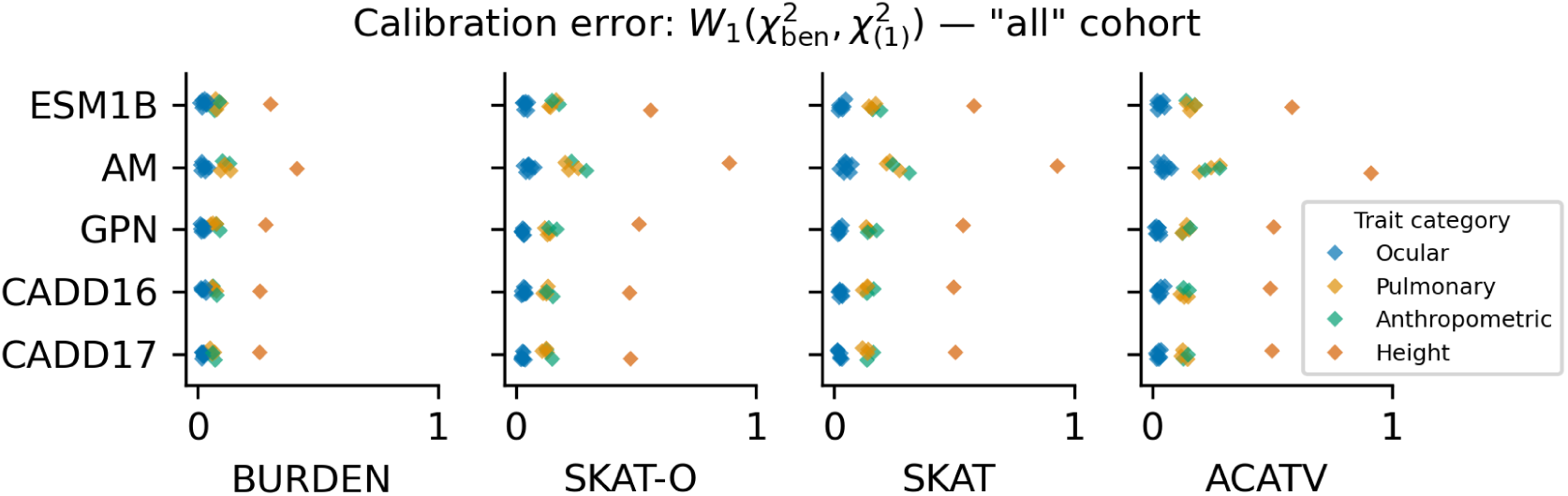
Null calibration by primary test and annotation method. Null calibration (*W*_1_ distance between benign-mask chi-squared statistics and the null 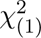 distribution) of primary test methods (subpanels) in the “all” cohort using variants classified as benign by each of the annotation methods (*y*-axis bins). Each point corresponds to one of 14 traits × 5 annotations × 4 tests = 280 RVATs, and is colored by phenotype category.

**Fig. S11:**
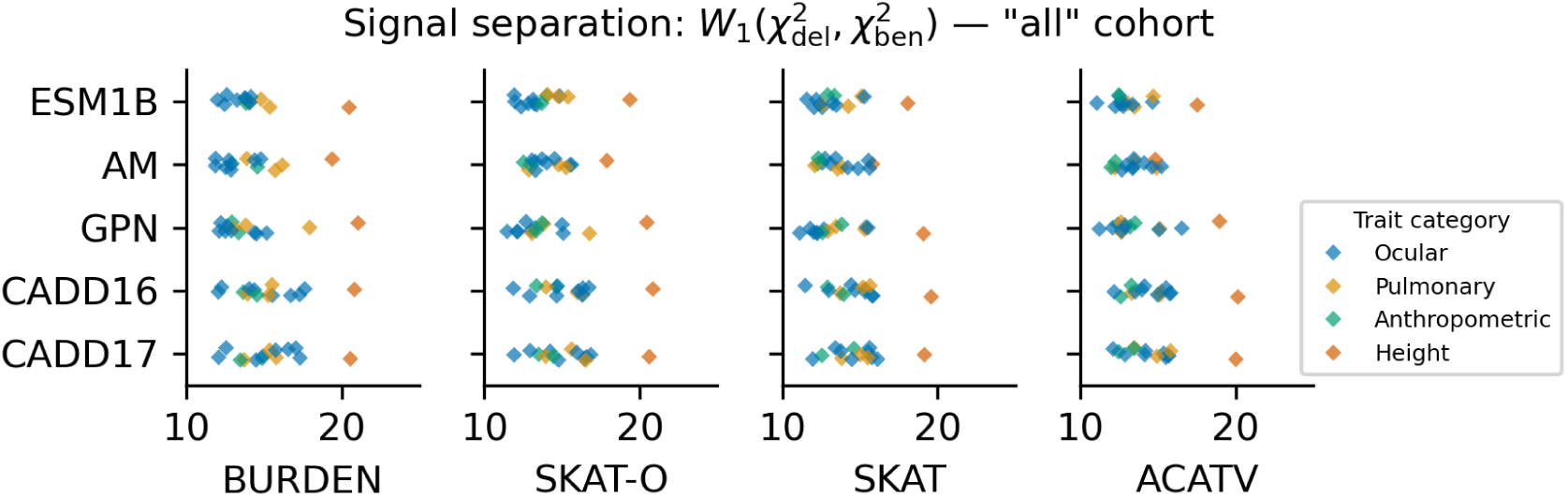
Signal separation by primary test and annotation method. Signal separation (*W*_1_ distance between benign-mask and deleterious-mask chi-squared statistics) of primary test methods (subpanels) in the “all” cohort using labels from each of the annotation methods (*y*-axis bins). Each point corresponds to one of 14 traits × 5 annotations × 4 tests = 280 RVATs, and is colored by phenotype category.

**Fig. S12:**
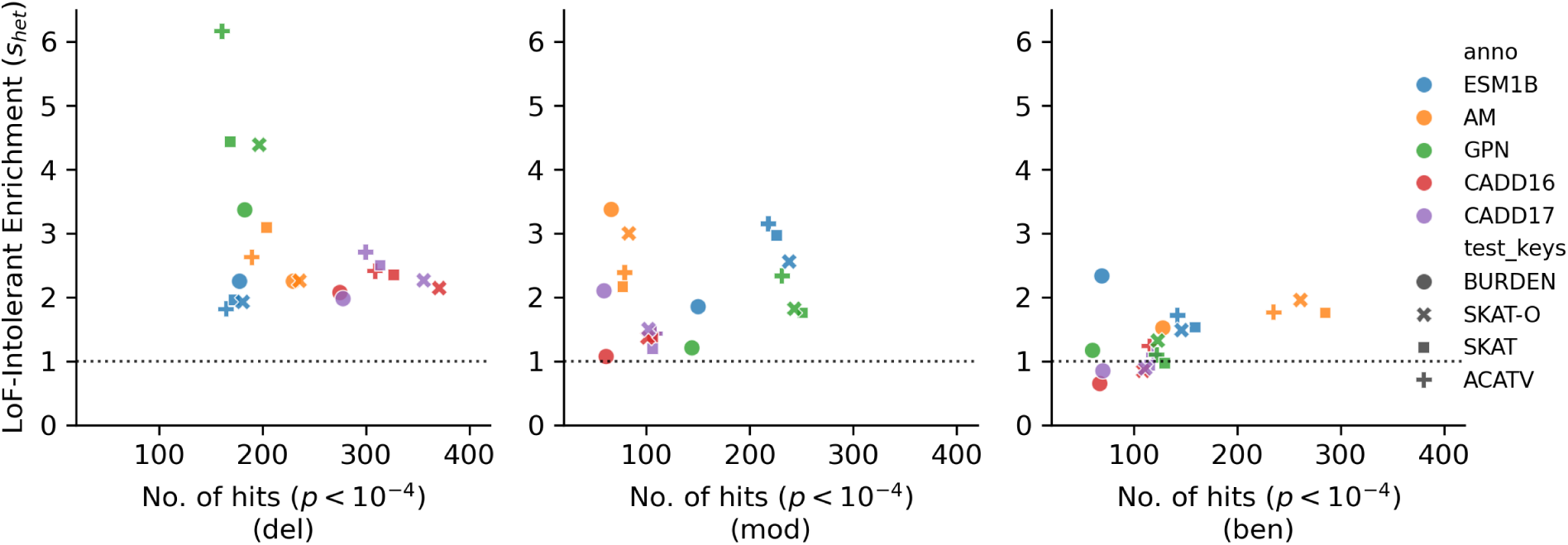
Signal enrichment in LoF-intolerant genes by variant mask for primary tests. The total number of hits for each primary association test (significant gene-trait pairs with *p <* 10*^−^*^4^ summed across all 14 traits; *x*-axis), and the enrichment of those genes in the top quartile (largest 25%) of *s*_het_ compared to the bottom quartile (*y*-axis). Each panel corresponds to a distinct set of input variants for each annotation method, defined by how it labels of variants as benign, moderate, or deleterious. In each panel, points are shaped by test and colored by annotation method.

**Fig. S13:**
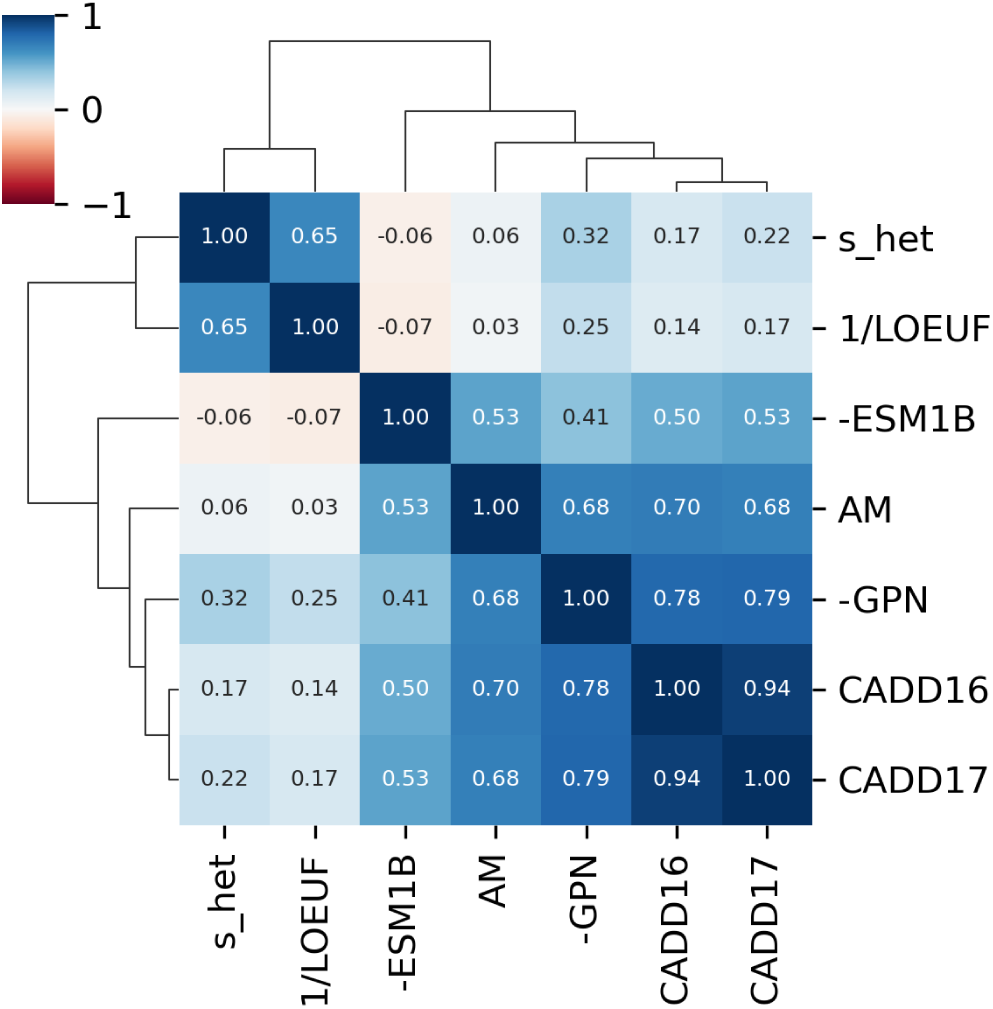
Clustered heatmap of raw deleteriousness scores from each annotation methods for missense variants, along with measures of gene-level intolerance to loss-of-function variation. Numbers in each box indicate the rank correlation (Spearman *ρ*) between pairs of methods. Scores from ESM and GPN are negated and LOEUF scores are inverted to polarize scores from all methods to the same direction — a larger score for a variant means it is more deleterious, and a larger score for a gene means that it is under stronger selection.

**Fig. S14:**
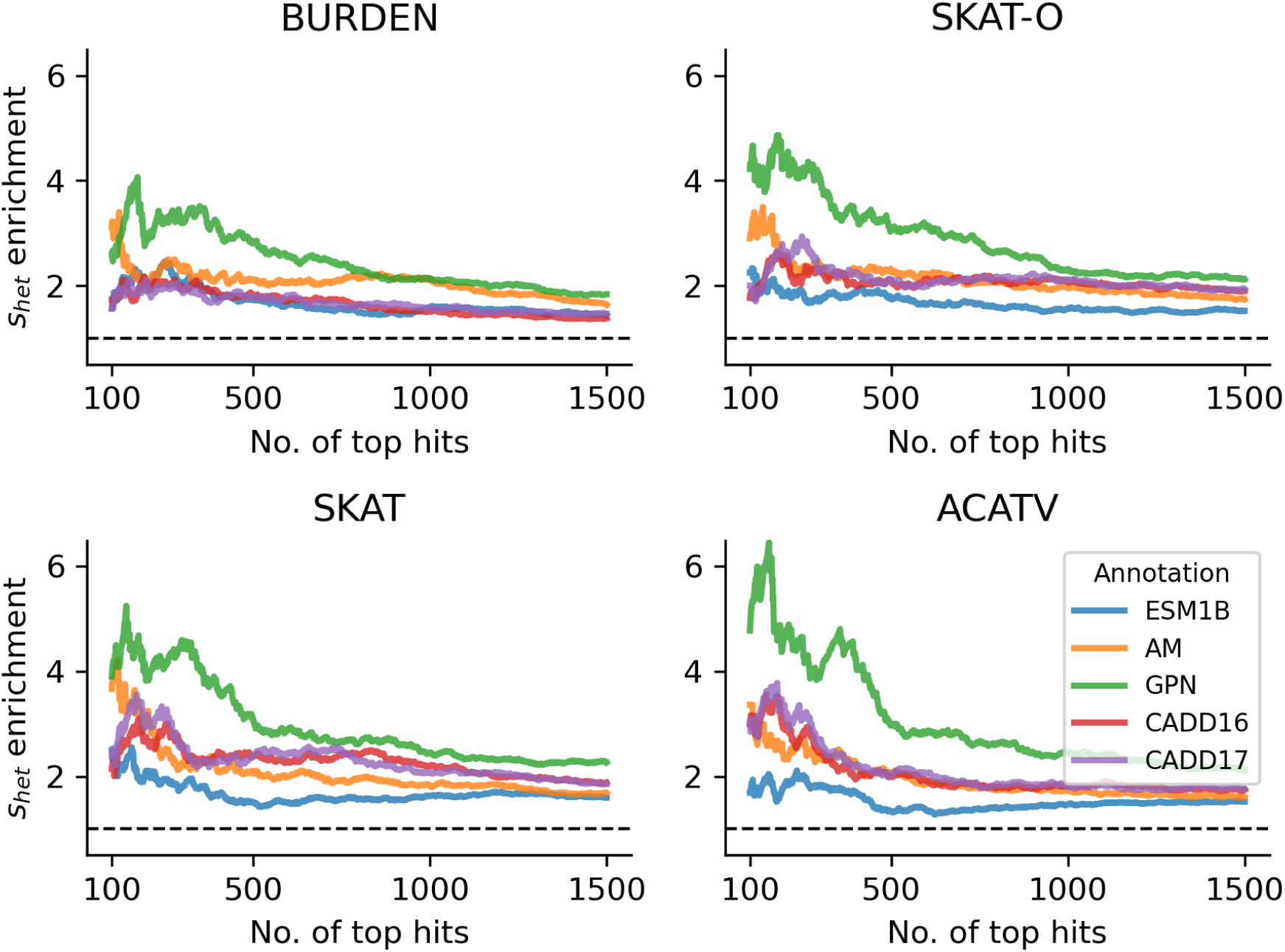
Signal enrichment in loss-of-function intolerant genes, as a function of top hits. For each primary test and annotation method, the enrichment of high-*s*_het_ genes (top quartile over bottom quartile) among the top *k* gene-trait pairs (ranked by *p*-value) nominated by the test using a deleterious variant mask from the annotation method. Colors are as in Fig. 4.

**Fig. S15:**
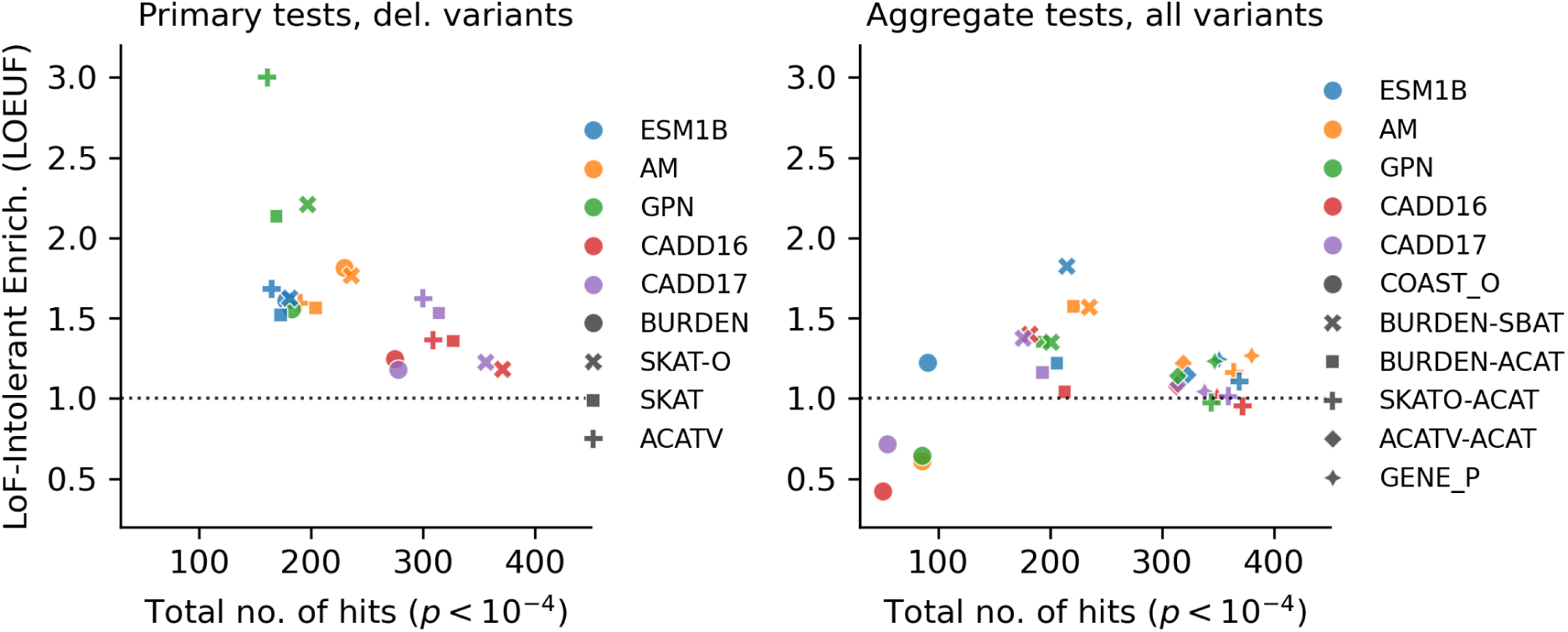
Signal enrichment in loss-of-function intolerant genes. For each primary (left panel) and secondary (right panel) test, the total number of hits (significant gene-trait pairs with *p <* 10*^−^*^4^ summed across all 14 traits), and the enrichment of those genes in the top quartile (largest 25%) of LOEUF compared to the bottom quartile. Points are shaped by test and colored by annotation method.

**Fig. S16:**
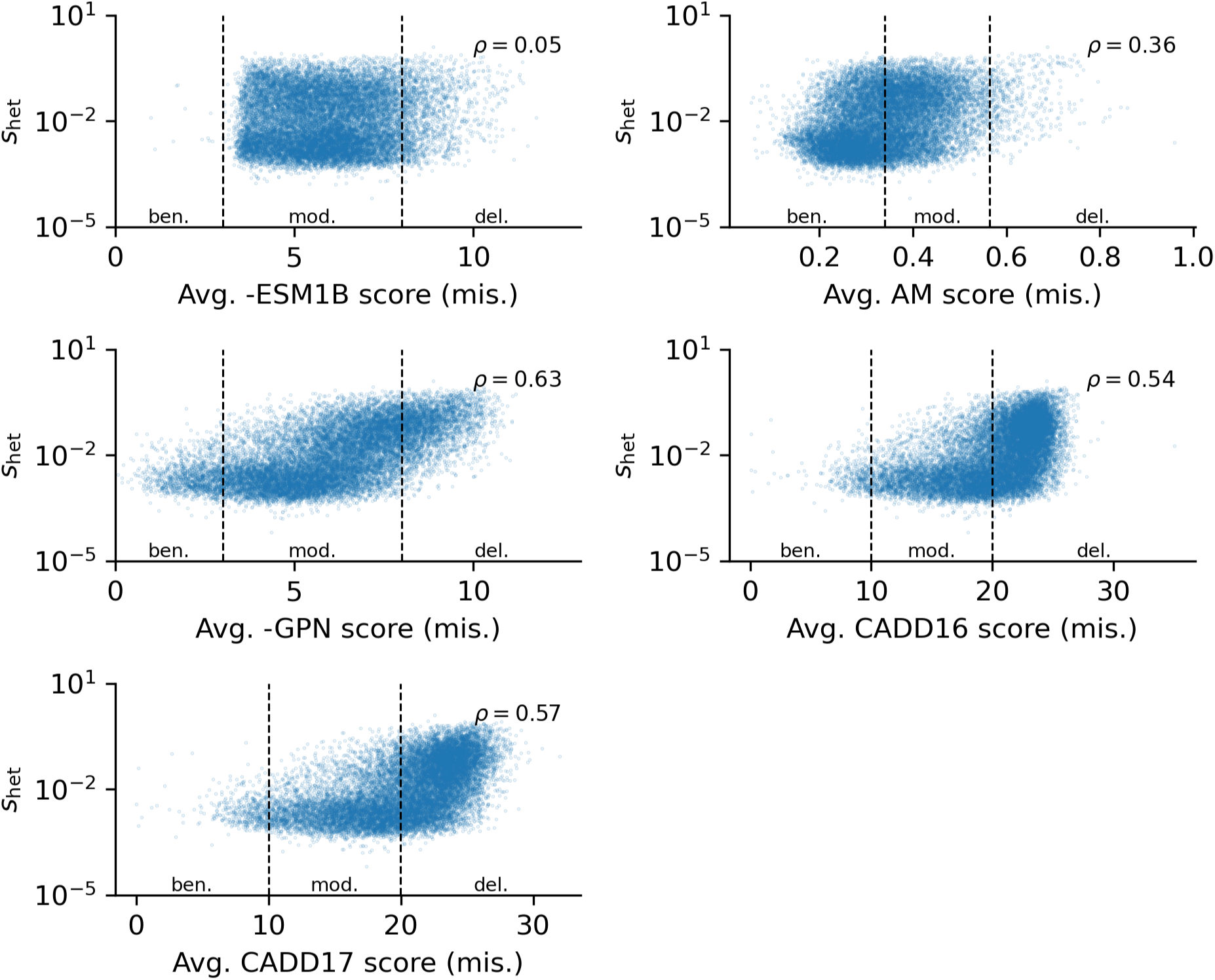
Relationship between selection against heterozygous loss-of-function variation (*s*_het_) and missense pathogenicity. For each annotation method (subpanels), scatterplots of the mean pathogenicity score of missense variants in each gene, and its estimated value of *s*_het_ [27]. Panels are annotated with the rank correlation between the mean missense pathogenicity score and *s*_het_, and with the method-specific thresholds used to define variants as benign, moderate, or pathogenic.

## References

[1] Wang, Q. et al. Rare variant contribution to human disease in 281,104 UK Biobank exomes. Na-ture 597, 527–532 (2021). URL https://www.nature.com/articles/s41586-021-03855-y.

[2] Backman, J. D. et al. Exome sequencing and analysis of 454,787 UK Biobank participants. Na-ture 599, 628–634 (2021). URL https://www.nature.com/articles/s41586-021-04103-z.

[3] Sun, B. B. et al. Genetic associations of protein-coding variants in human disease. Nature 603, 95–102 (2022). URL https://www.nature.com/articles/s41586-022-04394-w.

[4] Weiner, D. J. et al. Polygenic architecture of rare coding variation across 394,783 exomes. Na-ture 614, 492–499 (2023). URL https://www.nature.com/articles/s41586-022-05684-z.

[5] Spence, J. P. et al. Specificity, length and luck drive gene rankings in association studies. Nature 649, 918–925 (2026). URL https://www.nature.com/articles/s41586-025-09703-7.

[6] Schwartzentruber, J., Fiziev, P. P., McRae, J., Ulirsch, J. C. & Farh, K. K.-H. Flexibly Modeling Rare Variant Pathogenicity Improves Gene Discovery for Complex Traits (2025). URL https://www.biorxiv.org/content/10.1101/2025.11.04.686320v1. ISSN: 2692-8205 Pages: 2025.11.04.686320 Section: New Results.

[7] Han, S. et al. MIRAGE: A Bayesian statistical method for gene-level rare-variant analysis incorporating functional annotations. The American Journal of Human Genetics 113, 168–183 (2026). URL https://www.cell.com/ajhg/abstract/S0002-9297(25)00438-0.

[8] Rentzsch, P., Schubach, M., Shendure, J. & Kircher, M. CADD-Splice—improving genome-wide variant effect prediction using deep learning-derived splice scores. Genome Medicine 13, 31 (2021). URL 10.1186/s13073-021-00835-9.

[9] Schubach, M., Maass, T., Nazaretyan, L., Röner, S. & Kircher, M. CADD v1.7: using protein language models, regulatory CNNs and other nucleotide-level scores to improve genome-wide variant predictions. Nucleic Acids Research 52, D1143–D1154 (2024). URL 10.1093/nar/gkad989.

[10] Cheng, J., et al. Accurate proteome-wide missense variant effect prediction with AlphaMis-sense. Science 381, eadg7492 (2023). URL https://www.science.org/doi/full/10.1126/science.adg7492.

[11] Rives, A. et al. Biological structure and function emerge from scaling unsupervised learn-ing to 250 million protein sequences. Proceedings of the National Academy of Sciences 118, e2016239118 (2021). URL https://www.pnas.org/doi/10.1073/pnas.2016239118.

[12] Benegas, G., Albors, C., Aw, A. J., Ye, C. & Song, Y. S. A DNA language model based on multispecies alignment predicts the effects of genome-wide variants. Nature Biotechnology 43, 1960–1965 (2025). URL https://www.nature.com/articles/s41587-024-02511-w.

[13] Landrum, M. J. et al. ClinVar: public archive of interpretations of clinically relevant vari-ants. Nucleic Acids Research 44, D862–D868 (2016). URL 10.1093/nar/gkv1222.

[14] Brandes, N., Goldman, G., Wang, C. H., Ye, C. J. & Ntranos, V. Genome-wide prediction of disease variant effects with a deep protein language model. Nature Genetics 55, 1512–1522 (2023). URL https://www.nature.com/articles/s41588-023-01465-0.

[15] Chen, S. et al. A genomic mutational constraint map using variation in 76,156 human genomes. Nature 625, 92–100 (2024). URL https://www.nature.com/articles/s41586-023-06045-0.

[16] Bose, D., Fuchsberger, C. & Boehnke, M. Rare-variant association studies: When are ag-gregation tests more powerful than single-variant tests? The American Journal of Human Genetics 112, 1948–1961 (2025). URL https://www.sciencedirect.com/science/article/pii/S0002929725002721.

[17] Liu, Y. et al. ACAT: A Fast and Powerful p Value Combination Method for Rare-Variant Analysis in Sequencing Studies. The American Journal of Human Genetics 104, 410–421 (2019). URL https://www.cell.com/ajhg/abstract/S0002-9297(19)30002-3.

[18] Wu, M. C. et al. Rare-Variant Association Testing for Sequencing Data with the Sequence Kernel Association Test. The American Journal of Human Genetics 89, 82–93 (2011). URL https://www.cell.com/ajhg/abstract/S0002-9297(11)00222-9.

[19] Lee, S. et al. Optimal Unified Approach for Rare-Variant Association Testing with Applica-tion to Small-Sample Case-Control Whole-Exome Sequencing Studies. The American Jour-nal of Human Genetics 91, 224–237 (2012). URL https://www.cell.com/ajhg/abstract/S0002-9297(12)00316-3.

[20] Lee, S., Abecasis, G. R., Boehnke, M. & Lin, X. Rare-Variant Association Analysis: Study Designs and Statistical Tests. The American Journal of Human Genetics 95, 5–23 (2014). URL https://www.sciencedirect.com/science/article/pii/S0002929714002717.

[21] Ziyatdinov, A. et al. Joint testing of rare variant burden scores using non-negative least squares. The American Journal of Human Genetics 111, 2139–2149 (2024). URL https://www.sciencedirect.com/science/article/pii/S0002929724003070.

[22] McCaw, Z. R. et al. An allelic-series rare-variant association test for candidate-gene dis-covery. The American Journal of Human Genetics 110, 1330–1342 (2023). URL https://www.sciencedirect.com/science/article/pii/S0002929723002410.

[23] Mbatchou, J. et al. Computationally efficient whole-genome regression for quantitative and bi-nary traits. Nature Genetics 53, 1097–1103 (2021). URL https://www.nature.com/articles/s41588-021-00870-7.

[24] Bycroft, C. et al. The UK Biobank resource with deep phenotyping and genomic data. Nature 562, 203–209 (2018). URL https://www.nature.com/articles/s41586-018-0579-z.

[25] Devlin, B. & Roeder, K. Genomic Control for Association Studies. Biometrics 55, 997–1004 (1999). URL 10.1111/j.0006-341X.1999.00997.x.

[26] Karczewski, K. J. et al. Systematic single-variant and gene-based association testing of thou-sands of phenotypes in 394,841 UK Biobank exomes. Cell Genomics 2, 100168 (2022). URL https://www.sciencedirect.com/science/article/pii/S2666979X22001100.

[27] Zeng, T., Spence, J. P., Mostafavi, H. & Pritchard, J. K. Bayesian estimation of gene constraint from an evolutionary model with gene features. Nature Genetics 56, 1632–1643 (2024). URL https://www.nature.com/articles/s41588-024-01820-9.

[28] McCaw, Z. R. et al. Pitfalls in performing genome-wide association studies on ratio traits. Hu-man Genetics and Genomics Advances 6, 100406 (2025). URL https://www.sciencedirect.com/science/article/pii/S2666247725000090.

[29] Zhou, W. et al. Scalable generalized linear mixed model for region-based association tests in large biobanks and cohorts. Nature Genetics 52, 634–639 (2020). URL https://www.nature.com/articles/s41588-020-0621-6.

[30] Zhou, W. et al. SAIGE-GENE+ improves the efficiency and accuracy of set-based rare variant association tests. Nature Genetics 54, 1466–1469 (2022). URL https://www.nature.com/articles/s41588-022-01178-w.

[31] Guo, M. H. et al. Determinants of Power in Gene-Based Burden Testing for Monogenic Disorders. The American Journal of Human Genetics 99, 527–539 (2016). URL https://www.sciencedirect.com/science/article/pii/S0002929716302695.

[32] Jumper, J. et al. Highly accurate protein structure prediction with AlphaFold. Nature 596, 583–589 (2021). URL https://www.nature.com/articles/s41586-021-03819-2.

[33] Liu, Y. & Xie, J. Cauchy Combination Test: A Powerful Test With Analytic p-Value Calcula-tion Under Arbitrary Dependency Structures. Journal of the American Statistical Association 115, 393–402 (2020). URL https://doi.org/10.1080/01621459.2018.1554485. _eprint: 10.1080/01621459.2018.1554485.

[34] Karczewski, K. J. et al. The mutational constraint spectrum quantified from variation in 141,456 humans. Nature 581, 434–443 (2020). URL https://www.nature.com/articles/s41586-020-2308-7.

